# Metabarcoding and ecological interaction networks for selecting candidate biological control agents

**DOI:** 10.1101/2021.05.04.442560

**Authors:** Mélodie Ollivier, Vincent Lesieur, Johannes Tavoillot, Fanny Bénetière, Marie-Stéphane Tixier, Jean-François Martin

## Abstract

1. Classical biological control can be used to decrease the density of invasive species to below an acceptable ecological and economic threshold. Natural enemies specific to the invasive species are selected from its native range and released into the invaded range. This approach has drawbacks, despite the performance of specificity tests to ensure its safety, because the fundamental host range defined under controlled conditions does not represent the actual host range *in natura*, and these tests omit indirect interactions within community.
2. We focus on *Sonchus oleraceus* (Asteraceae), a weed species originating from Western Palearctic that is invasive worldwide and notably in Australia. We explore how analyses of interaction network within its native range can be used to 1) inventory herbivores associated to the target plant, 2) characterize their ecological host ranges, and 3) guide the selection of candidate biocontrol agents considering interactions with species from higher trophic levels. Arthropods were collected from plant community sympatric to *S. oleraceus*, in three bioclimatic regions, and interactions were inferred by a combination of molecular and morphological approaches.
3. The networks reconstructed were structured in several trophic levels from basal species (diversified plant community), to intermediate and top species (herbivorous arthropods and their natural enemies). The subnetwork centered on *S. oleraceus* related interactions contained 116 taxa and 213 interactions. We identified 47 herbivores feeding on *S. oleraceus*, 15 of which were specific to the target species (*i.e*. Generality value equal to 1). Some discrepancies with respect to published findings or conventional specificity tests suggested possible insufficient sampling effort for the recording of interactions or the existence of cryptic species. Among potential candidate agents, 6 exhibited interactions with natural enemies.
4. *Synthesis and applications:* Adopting a network approach as prerequisite step of the CBC program can provide a rapid screening of potential agents to be tested in priority. Once ecological host range defined, we suggest that priority should be given to agent predated by a minimum species, and, when they exist, to an agent that possesses enemies from the most distant taxonomical group from those occurring in the range of introduction.

## Introduction

Biological invasions are currently threatening biodiversity to an unprecedented extent (Bellard et al., 2016; Seebens et al., 2015; Vitousek et al., 1997). When invasive species disrupt the ecological or economic balance, action is required to control their negative impact. Chemical control methods are widely used in such situations, but classical biological control (CBC) constitutes a possible alternative. CBC involves the release of natural enemies, specific to the target organism and originating from its native range, to keep the density of the invasive species below an economically and ecologically acceptable threshold (Keane and Crawley, 2002; McFadyen, 1998; Van Driesche et al., 2010). CBC is considered more sustainable than chemical control (Peterson et al., 2020), although the introduction of biocontrol agents (BCA) into a new territory may itself represents a risk for the recipient communities (Barratt et al., 2018; Hinz et al., 2019; Suckling and Sforza, 2014). Once introduced, the BCA may affect non-target species, especially if it lacks specificity (Müller-Schärer and Schaffner, 2008). Assessing the host range of a candidate BCA is, thus, crucial, to anticipate such risks. Most of the host-specificity tests performed to assess this risk are conducted under standardized conditions, through choice/no-choice experiments over a range of targets selected according to the centrifugal phylogeny approach (Briese, 2005; Wapshere et al., 1989).

Recent reviews recognized the success of such experiments for limiting the undesirable unintentional effects of the CBC of weeds (Hinz et al., 2019, 2020). However, the cumbersome nature of these tests reduces the range of species that can be screened. The candidate BCA are selected through preliminary field monitoring that may miss a species of interest. Furthermore, as the fundamental host range of a species (defined under controlled conditions) is thought to be broader than the host range actually observed in the field (known as the realized host range; Louda et al., 2003; Schoonhoven et al., 1998; Sheppard and Harwood, 2005), these tests tend to overestimate the risk and lead to the rejection of candidate BCA based on interactions that would not occur in the field (false positives) (e.g. Groenteman et al. 2011). Most CBC programs use specificity tests under controlled conditions as proxies for field conditions due to the complexity of trophic interaction assessments in the field but this leaves room for improvement.

The characterization of ecological interactions among communities of plants and arthropods *in natura* is challenging, as it traditionally requires direct observations, the rearing of specimens and considerable taxonomic expertise, rendering the process impractical for large-scale studies. Recent advances in molecular approaches, such as the combination of DNA metabarcoding on gut content or feces and high-throughput next-generation sequencing (NGS), have opened up new opportunities to track the host range of arthropods *in natura* with both a high taxonomic resolution and high sensitivity (Derocles et al., 2018; Frei et al., 2019; Wirta et al., 2014; Zhu et al., 2019). Even interactions that are very difficult to observe, such as host-parasitoid associations, can be detected by such methods (Gariepy et al., 2014; Hrček and Godfray, 2015). This approach can be used to reconstruct networks of trophic interactions directly from studies in the field, and provides an analytical framework particularly relevant to studies of complex species assemblages. Network ecology do not only depicts species interactions, but provides elements for the understanding of recurrent patterns of antagonistic interactions between plants and herbivores, such as specialization or compartmentation (Lewinsohn et al., 2006; Thébault and Fontaine, 2010). In CBC against invasive weeds, analyses of ecological networks have been used to assess the extent to which a BCA fits into a recipient community. Such methods provided a way to quantify the direct impact of biological control on non-target plants (Memmott, 2000), and its indirect impact on other species at higher trophic levels (Carvalheiro et al., 2008; Louda et al., 1997; Pearson and Callaway, 2003). Such studies have highlighted the usefulness of network ecology for evaluating the impact of BCA after their introduction (Memmott, 2009; Willis and Memmott, 2005), but interaction network analysis can also be used for the upstream assessment of potential candidate BCA, in a more systematic process (Ollivier et al., 2020). Adopting a network approach as prerequisite step of the CBC program, can provide a rapid screening of the ecological host range of potential agents to be tested in priority. This can also inform about species functional properties through the position and connexions the species have in the network, independently of its taxonomic assignation, which would confer a strong predictive power of the interactions possibly occurring in a novel bioclimatic region (Todd et al., 2020). Indeed, the choice of BCA should also take into account indirect effects on the recipient community due to interactions with higher trophic levels in the network, *i.e*. natural enemies (Hinz et al., 2019; Memmott, 2000). If comparable enemies than those identified in the native range are present in the range of introduction, new interactions might be created with BCA, resulting in disturbances in the ecological network through indirect interactions, *e.g*. apparent competition (Carvalheiro et al., 2008; López-Núñez et al., 2017).

The objective of this study was to determine how the analysis of interaction networks could be used to support the selection of candidate BCA for the common sowthistle, *Sonchus oleraceus* L. (Asteraceae). This plant is native to Western Europe and Northern Africa (Boulos, 1974; Hutchinson et al., 1984) and is the most widely naturalized terrestrial plant worldwide (Pyšek et al., 2017). In Australia, it has become a weed of major concern in cropping systems (Llewellyn et al., 2016; Widderick et al., 2010). Aside the development of resistance to multiple herbicides (Adkins et al., 1997; Jalaludin et al., 2018; Meulen et al., 2016), the control of this weed is complex as it is extremely prolific and seeds can germinate all year round when sufficient rainfalls occurs. *Sonchus oleraceus* rapidly dominates crops, reducing yield and contaminating harvested grain (Llewellyn et al., 2016). A CBC program was therefore initiated in 2017, to identify candidate BCA. In this context, an analysis of ecological networks, based on direct field observations and high-throughput DNA metabarcoding, was performed. Our objectives were to 1) establish an inventory of arthropods feeding on *S. oleraceus*, and assess the contribution of the approach relatively to classical procedures, 2) delineate the ecological host range for herbivores feeding on *S. oleraceus* and point out candidate BCA, and 3) identify the trophic interactions of the candidate BCA with natural enemies, and consider their implications for the CBC program.

## Materials and Methods

### Sampling design

We maximized the species diversity and associated interactions, through a maximum variation design with three bioclimatic regions in France (semi-oceanic, Mediterranean and continental climates) (Ceglar et al., 2019) and three successive sampling dates (April, May and June 2018). Sampling was carried out from 10 A.M. to 4 P.M., by varying climatic conditions (wet, cloudy to sunny weathers and temperatures ranged between 10°C and 29°C). These variations did not affect our ability to capture arthropods. For each bioclimatic region and date, we employed an opportunistic sampling strategy to collect plants from several ruderal and agricultural sites, covering the diversity of habitats (open and disturbed) colonized by *S. oleraceus* (Supplementary Table 1). At each site, on each date, we sampled three quadrats (1 m^2^) along a 20 m linear transect. Quadrats were placed to contain at least one *S. oleraceus* plant. Within each quadrat, arthropods were collected from plants with a forceps or brush, and stored individually in sterile 2 ml Eppendorf tubes filled with a protective buffer solution. This solution is used to prevent oxidation of polyphenols and polyamines (PCR inhibitors) (see Cruaud et al. (2018) for more details). This procedure was repeated for each plant of every plant species present in the quadrat over a period of one hour, to standardize the sampling effort. This period was deemed adapted to represent the biodiversity of the sampled unit, and to allow vagrant insects, potentially disturbed by our arrival, to settle back on their resource plant before sampling. We collected individual specimens except for colonies of aphids, thrips, and egg masses, for which at least five specimens were required to obtain sufficient DNA for analysis. We did not consider pollinators or the soil fauna in this study. Following the collection of each specimen, tools were thoroughly cleaned by successive immersions in 2.5 % bleach solution, water and 96% ethanol, to prevent cross-contamination. At the end of the one-hour insect sampling period, all the plants within the quadrat were collected individually (by cutting the stem at the soil surface), for further dissection. Back in the laboratory, the plants were identified morphologically, and their organs (stems, leaves, flowers) were dissected to collect endophagous arthropods, which were transferred into tubes as described for the arthropods collected in the field. For each arthropod specimen collected, we identified the plant species from which arthropods were sampled, and recorded the specimen stage and condition (degraded, parasitized), and putative identification (at least taxonomic group, with identification to species level if straightforward). All arthropod samples were frozen at −20°C until DNA analysis. Thus, while plants were identified morphologically, arthropods were identified via molecular technologies. Each plant was transferred to a paper bag and oven-dried at 70°C for 72 h, for the determination of aboveground dry biomass (g) as an estimate of plant abundance per quadrat. Arthropod abundances were determined based on the number of individuals collected per quadrat for each taxon. Sampling was performed for 57 quadrats, over the three sampling dates.

### High-throughput DNA metabarcoding

We characterized the interaction network by directly observing plant-arthropod interactions (recording only interactions for which an observation of feeding was verified); while arthropod-arthropod interactions were revealed by molecular analysis. We first isolated total DNA from each arthropod individual (Cruaud et al., 2018). As presented in Supplementary Figure S1, we then performed metabarcoding on each arthropod sample, with a two-step DNA amplification and high-throughput sequencing method adapted from the procedure described by Galan et al. (2017). We sequenced three short COI fragments, with primer combinations and PCR protocols developed elsewhere (HCO forward: Leray et al. 2013, HCO reverse: Folmer et al. 1994, LEP F. and R.: Brandon-Mong et al. 2015, HEX F. and R.: Marquina et al. 2019), to overcome the problem of the lack of primer universality among arthropods. Error-proof indices for individual sample identification were developed with the high-throughput sequencing process described by Martin (2019). The libraries were sequenced with Illumina technology, using a Miseq 2×250 run for date 1 (April), and one lane of Hiseq 3000 each for dates 2 (May) and 3 (June).

The markers for each sample were demultiplexed with CutAdapt v2.3, and all paired-end reads were filtered for minimal length (280 bp), corrected for sequencing errors, and pairs of overlapping reads were merged with the Dada2 v1.12 R package (Callahan et al., 2016). A matrix was thus obtained, containing samples as variables and amplicon variant sequences (ASVS) as observations. A variant is a set of identical corrected and merged paired-end reads. We used Qiime2 (Bolyen et al., 2018) with a 2% divergence threshold, to merge ASVS, to decrease their number without the loss of taxonomic information. The summed number of reads for each merged variant for a given arthropod sample was reported as the intersection of samples and ASVS.

Each ASVS was assigned, by BLAST, to a barcoding reference database of cytochrome oxidase subunit I (COI) nucleotide sequences (658 bp) compiled from three different sources and curated by expert analysis. These reference barcodes were retrieved from BOLDSYSTEM (Ratnasingham and Hebert, 2007), the CBGP - Continental Arthropod collection (Centre de Biologie pour la Gestion des Population, 2019) and a local database specifically designed for this study. Our database contained barcodes of the most frequently encountered species during this sampling campaign (extra-specimens collected) and field surveys (2017-2020) conducted through Europe and North Africa for the search of *S. oleraceus* natural enemies (see below). In total, these three sources compiled 1 699 995 sequences from 119 299 species available for ASVS assignation. We retained successful assignments to the ranks of species, genus and family, but not those to higher taxonomic levels, because arthropod biology is too variable at higher taxonomic ranks to be informative for our purpose. The assignments obtained for each marker were grouped together in a single table and the numbers of reads were summed by assigned taxon. The resulting file was therefore an interaction matrix in BIOM file format, in which the assigned taxa replaced ASVS. The matrix was curated and manually transformed to obtain an adjacency matrix (in which the observations are sources and the variables are consumers) usable for further network analyses. For each pair of consumer/prey species, occurrence frequencies of interaction were calculated (Supplementary Text 1 and Figure S2).

#### Assessment of sampling robustness and global network description

We first evaluated the completeness of sampling over the entire sampling campaign, and generated taxon accumulation curves (the 57 quadrats were added in a random order, with 1,000 permutations) for plants and arthropods, using the *specaccum* function of the R package *vegan* (Oksanen *et al*., 2019). We estimated the extrapolated taxonomic richness by calculating the Chao 1 index (Chao, 1984) with the *specpool* function. Likewise, the robustness of sampling for the characterization of interactions was assessed by generating accumulation curves for pairwise interactions. We first generated an accumulation curve including all the types of direct interactions (*e.g*. plant-herbivores, herbivores-natural enemies, etc.) present in the meta-network (*i.e*. pooling interactions from all sites). The 57 quadrats were added in a random order, with 1000 permutations. We finally generated a curve focusing on interactions involving *S. oleraceus* as a source, to evaluate the performance of the sampling design for addressing our objective of establishing an inventory of the arthropods feeding on *S. oleraceus*, corresponding to candidate BCA. For both curves, we estimated the extrapolated interaction richness with the Chao 1 index (Chao 1984), using the *specpool* function.

Prior to interaction analyses, a global description of the meta-network (pooling interactions data from all sites) and subnetwork (centred on *S. oleraceus* related interactions) was performed. Several metrics were calculated: the number of links (L), the number of nodes (S) (connected and isolated), connectance (C) and link density (LD) (Bersier et al., 2002; Warren, 1994). Connectance is the proportion of the possible trophic links actually realized; here cannibalism is not permitted, so C = L/S(S-1). Link density is the mean number of links per taxon, calculated as LD = L/S. We also characterized the taxon assemblage by determining taxonomic richness (*i.e*. number of taxa) for each trophic level (plants, herbivores and natural enemies).

#### Selection of candidate biocontrol agents

The selection of candidate BCA was decided according two criteria: a restricted ecological host range and limited interactions with natural enemies. Thus, based on the interactions retrieved from the meta-network, we selected a subnetwork considering only the arthropods having *S. oleraceus* as a source plant, as well as all their complementary plant resources. We also included natural enemies associated with these herbivores (*i.e*. parasitoids and predators). We assessed and visualized the specificity of these herbivores, by plotting interactions between herbivores encountered on *S. oleraceus* and all their complementary resource plants as a grid matrix, in which plants were ordered by their degree of phylogenetic relatedness to *S. oleraceus*, as defined by the current classification of angiosperms (Chase et al., 2016). Arthropods were ordered by the increasing generality values (*i.e*. the number of resources per taxon) characterizing ecological host range. To assess and visualize the dependence of natural enemies on these herbivores, we constructed a second level grid matrix in connexion with the previous, and calculated arthropod vulnerability values (*i.e*. the number of consumer per taxon). Multipartite network and grid matrices were constructed with *igraph* R package.

#### Assessing the contribution of the approach for the biocontrol program

To discuss the contribution of the method herein proposed, we used, as a point of reference, a survey performed following classical procedures (sampling, rearing and identification of specimens exclusively collected from *S. oleraceus*) in the frame of this CBC program (Lesieur et al., in prep). However, we acknowledge that this classical survey covered a longer period of sampling (2017-2020) and a much larger geographical area was prospected (10 countries through Europe and North Africa).

## Results

### Summary of the molecular results

In total, 2,834 arthropod specimens were collected and analyzed by metabarcoding, to reconstruct the interaction network at a global scale. We obtained DNA sequences and taxonomic assignments for 1,803 of the 2,834 arthropods initially collected (63.6 %). This proportion of exploitable information reached 71% (2,011 specimens) after manual validation of the matrix.

The molecular analysis provided a total of 107,483,410 reads, 19.2% of which were retained after screening with quality filters; we obtained a final dataset of 2,014 COI variant sequences (Supplementary Table 2). Before, manual validation, we observed that a large proportion of the diversity (33% of the families and 40% of the species) was recovered by the use of all markers, the rest being recovered by a combination of two markers, or specifically found with only one marker (Supplementary Figure S3). LEP increased identification rates by 20% for families and 25% for total species, consistent with its widespread use in the research community (Brandon-Mong et al., 2015). The other two markers also provided original information, albeit to a lesser extent, at least as far as the number of taxa recovered was concerned, as 15% of the families and 16% of the species would not have been recovered with LEP alone. After data validation, 269 taxa were identified for arthropods, with 84% identified to species level (17 orders, 90 families and 189 genera). While plant taxonomic diversity (relying on morphological identifications) accounted for 132 taxa, 80% of which were classified to species level (25 orders, 29 families and 87 genera) (Supplementary figure S4).

#### Sampling robustness

The accumulation curve of plants seemed to approach an asymptote, but this was not the case for arthropods (Figure 1). The Chao 1 index indicated an extrapolated taxonomic richness value for plants of 164 taxa (± 12), with 132 taxa actually sampled. By contrast, for arthropods, the extrapolated taxonomic richness value was 442 taxa (± 39), but only 269 taxa were actually sampled. Sampling robustness was high over the entire sampling scheme for plants but sampling efficiency was lower for arthropods. Likewise, we assessed the completeness of pairwise interactions detected over the whole network. We observed a linear increase associated with a Chao1 index of 1245 (± 183) expected interactions, where 350 links were actually reconstructed (Figure 1). However, this is less of an issue for interactions involving *S. oleraceus*, the focus of the analysis for which this sampling was designed. The accumulation curve in question tended towards an asymptote, with a Chao 1 index of 63 (± 10) expected interactions and 47 interactions sampled. Overall, these results suggest that the sampling effort was adequate for the reconstruction of a unique interaction network maximizing of the proportion of links observed (Jordano, 2016).

**Figure 1:**
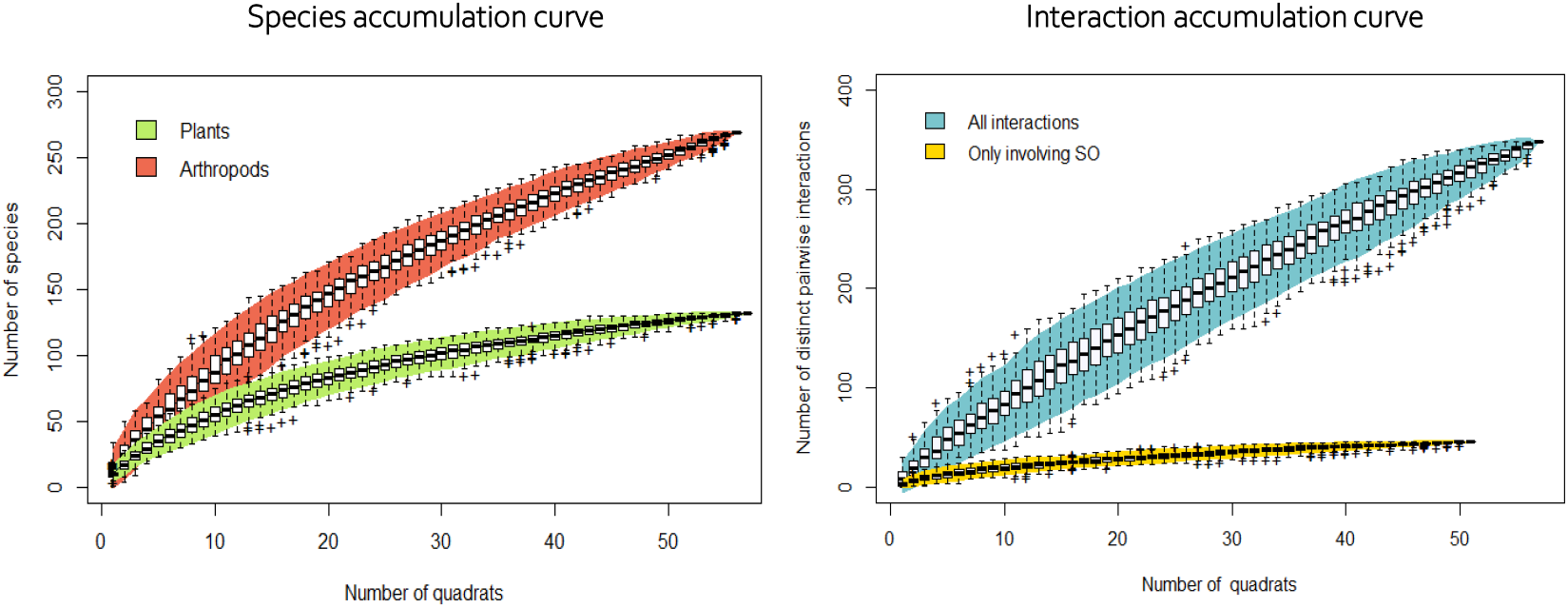
Accumulation curves representing a) species richness in plants and arthropods and b) pairwise interactions from the meta-network and focusing on interactions involving *S. oleraceus*. Curves were constructed with 1000 random resampling events over the 57 quadrats analyzed along the sampling campaign (Spring 2018).

#### Meta-network and subnetwork analyses

As presented in Table 1, the complete interaction network (meta-network) consisted of 401 nodes, 241 of which were connected to another node (60%), resulting in 350 links (Supplementary Figure S5). Linkage density and connectance calculated were 1.45 and 0.006, respectively. The meta-network included 60 plants in interaction (46% of the plants collected), 136 herbivores in interaction (74% of the herbivores collected), 35 natural enemies in interaction (49% of the natural enemies collected) comprising 19 parasitoid and 16 predator taxa, and 10 omnivores (feeding at more than one trophic level). The sub-network consisted of 116 nodes and 213 links, and resulting linkage density and connectance were 1.84 and 0.008, respectively (Figure 2). A more detailed description of taxon assemblage composing *S. oleraceus* subnetwork is provided in the following section.

**Figure 2:**
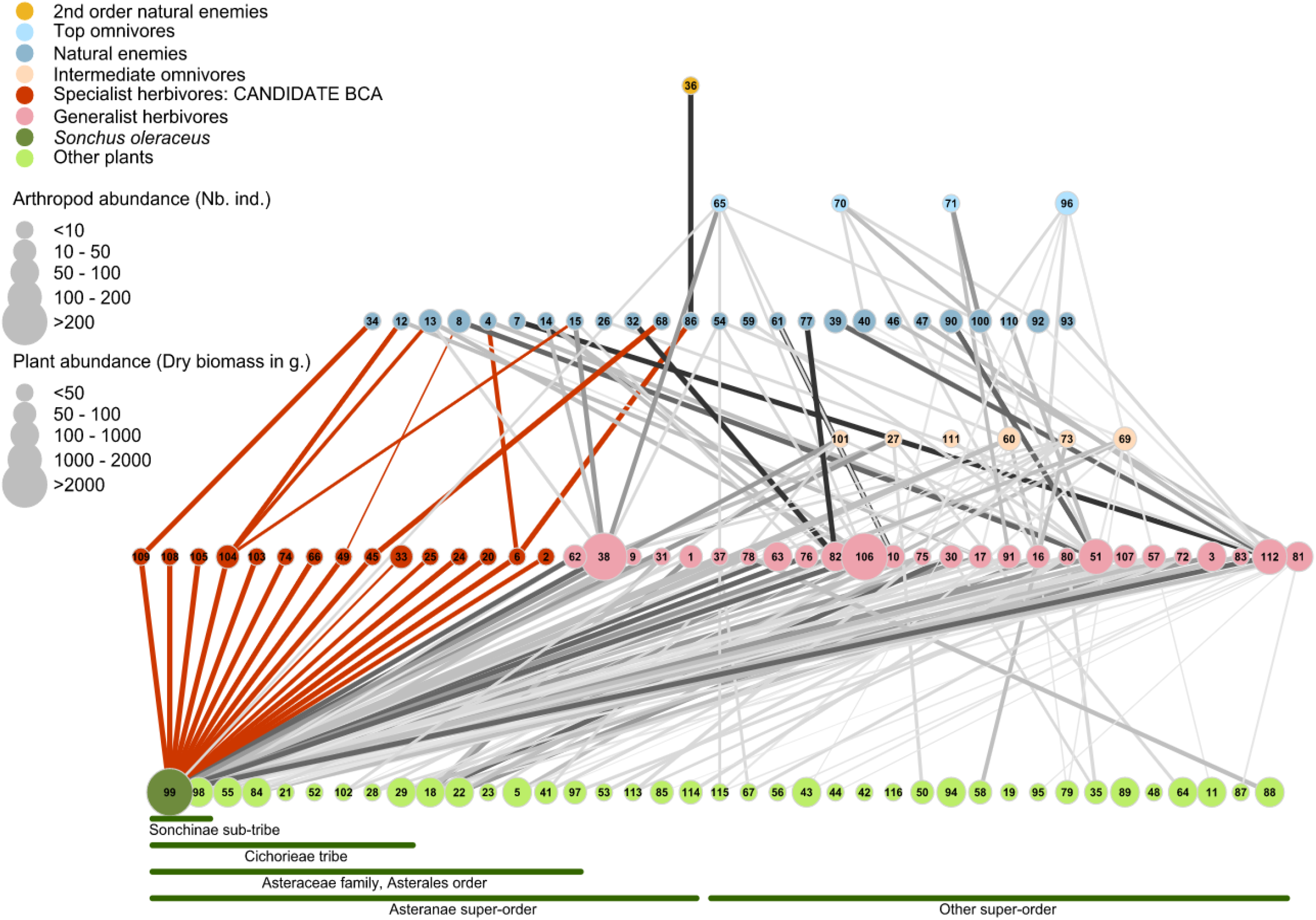
Multitrophic network reconstructed from *S. oleraceus* (dark green node) and 38 other plants (light green nodes) used by *S. oleraceus* herbivores (red nodes represent putative specialist herbivores, and pink nodes correspond to herbivores feeding on other species in addition to S. oleraceus). Plants are ordered by their phylogenetic relatedness to S. oleraceus. Natural enemies of herbivore species are represented by dark blue nodes. Nodes at intermediate levels (beige and light blue) correspond to omnivorous species identified by molecular analyses. The width of edges reflects frequencies of interactions between pairs of species and edges coloured in red emphasize interactions involving potential candidate BCA. The list of taxa corresponding to each node and the edge list are provided in Supplementary Tables 3 and 4, respectively.

**Table 1:** Global description of the meta-network and subnetwork centred on *S. oleraceus*. Metrics measured are the total number of nodes (in brackets is the number of nodes connected in the network), the number of links, the linkage density and the connectance of the network. For each network is also presented the number of species per trophic level (in brackets is the number of species in interaction with another species). Omnivorous species are regarded as natural enemies.

|  |  | Meta-network | Subnetwork* |
| --- | --- | --- | --- |
| **Global metrics** | Nb of nodes (connected) | 401 (241) | 116 |
|  | Nb of links | 350 | 213 |
|  | Linkage density | 1,45 | 1,84 |
|  | Connectance | 0,006 | 0,008 |
| **Taxon assemblage** | Nb of plant species (in interaction) | 132 (60) | 39 |
|  | Nb of herbivore species (in interaction) | 185 (136) | 47 |
|  | Nb of natural enemies species (in interaction) | 79 (35) | 30 |

\* All nodes are connected

Identifying candidate biocontrol agents: considering host range and regulation by enemies Analysing *S. oleraceus* subnetwork, we found 47 herbivorous taxa feeding on the target, including 37 taxa identified to species level. They belonged to five different orders, *i.e*. Hemiptera (45%), Diptera (25%), Coleoptera (19%), Lepidoptera (0.06%) and Hymenoptera (0.04%), and were distributed in nine different trophic guilds, with the flower bud sucking-piercing guild being the most represented (23%) while the less represented guild corresponded to the chewing guild (2%) (Table 2). Fifteen taxa were collected exclusively from *S. oleraceus*, and another two taxa were collected from *S. oleraceus* and *Sonchus asper* (Figure 3). These taxa are potential BCA (host range apparently restricted to the genus *Sonchus*, subtribe Sonchinae). Six additional species were detected only on members of the tribe Chicorieae (*Aphis craccivora* Koch, *Ophiomyia cunctata* Hendel, *Phytomyza lateralis* Fallén, *Campiglosa producta* Loew, *L. punctiventris* and *T. formosa*). We identified 38 other plant species as complementary resource plants for the herbivore species collected from *S. oleraceus*. The generality of these herbivore species ranged from 1 to 18, with *Philaenus spumarius* L. the most polyphagous of the 47 herbivores species found on *S. oleraceus* (Figure 3).

**Figure 3:**
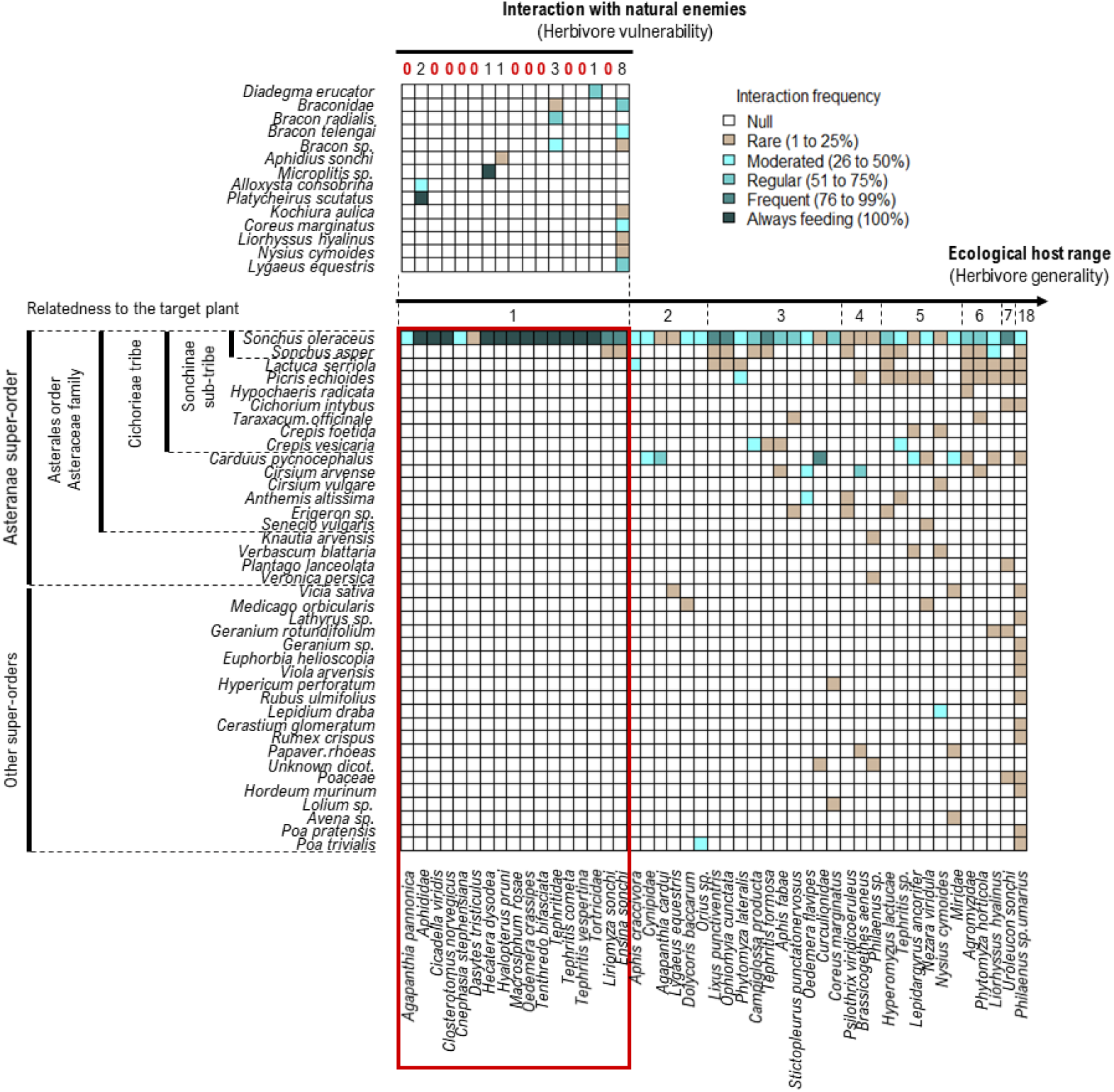
Interaction matrix between herbivores sampled on *Sonchus oleraceus* and their resource plants, indicating the ecological host range of the herbivores, as defined by intense field sampling in France, during spring 2018. Plants are ordered by their phylogenetic relatedness to *S. oleraceus* and arthropods are ordered by increasing generality values (i.e. the number of resources per species). Red rectangle highlights the 17 species considered as candidate BCA for their restricted eclogical host range. The second level matrix diplays interactions between candidate BCA and their natural enemies. Interaction are represented as semi-quantitative information, via occurences frequencies of interactions culculated for each species pair.

**Table 2:**
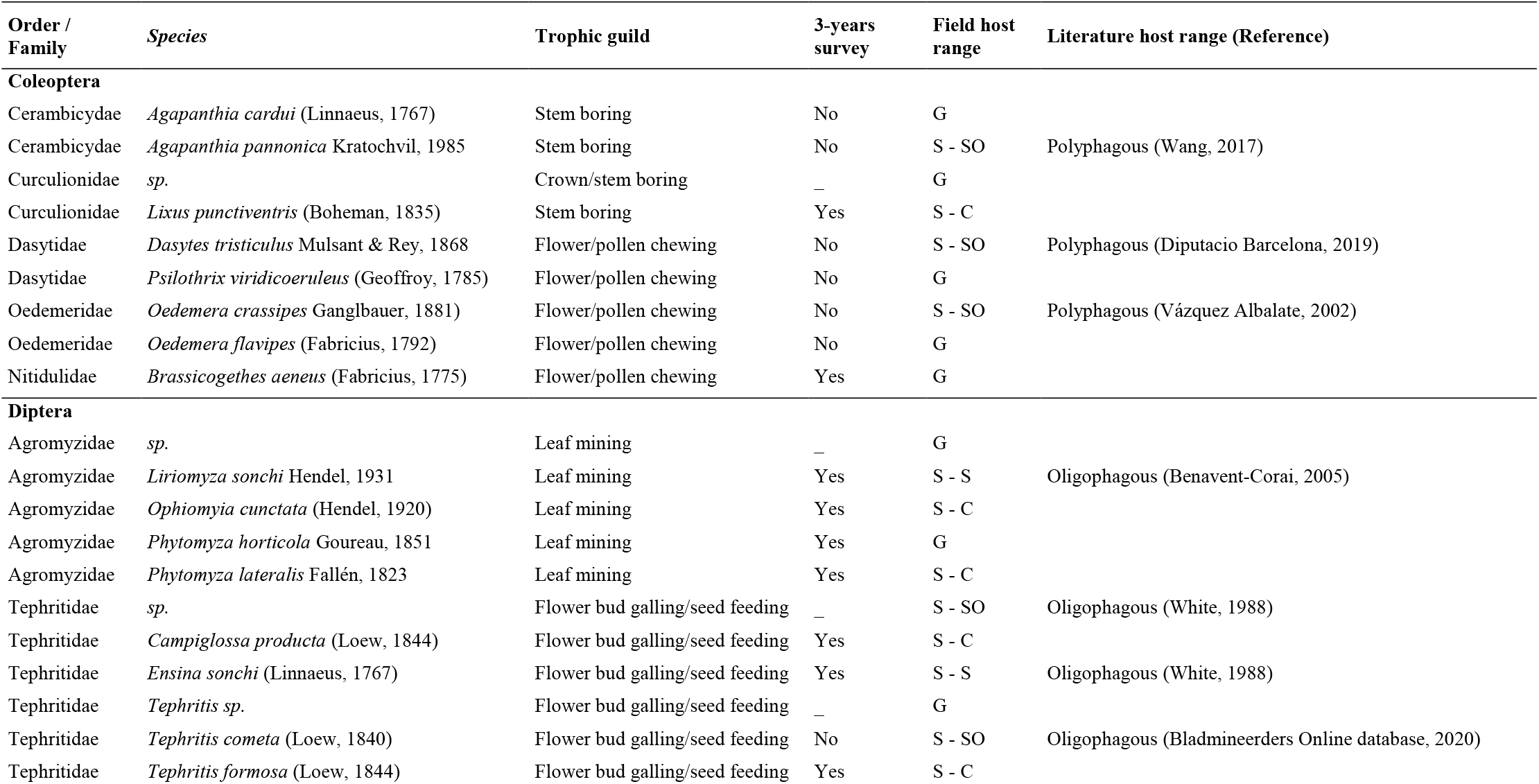
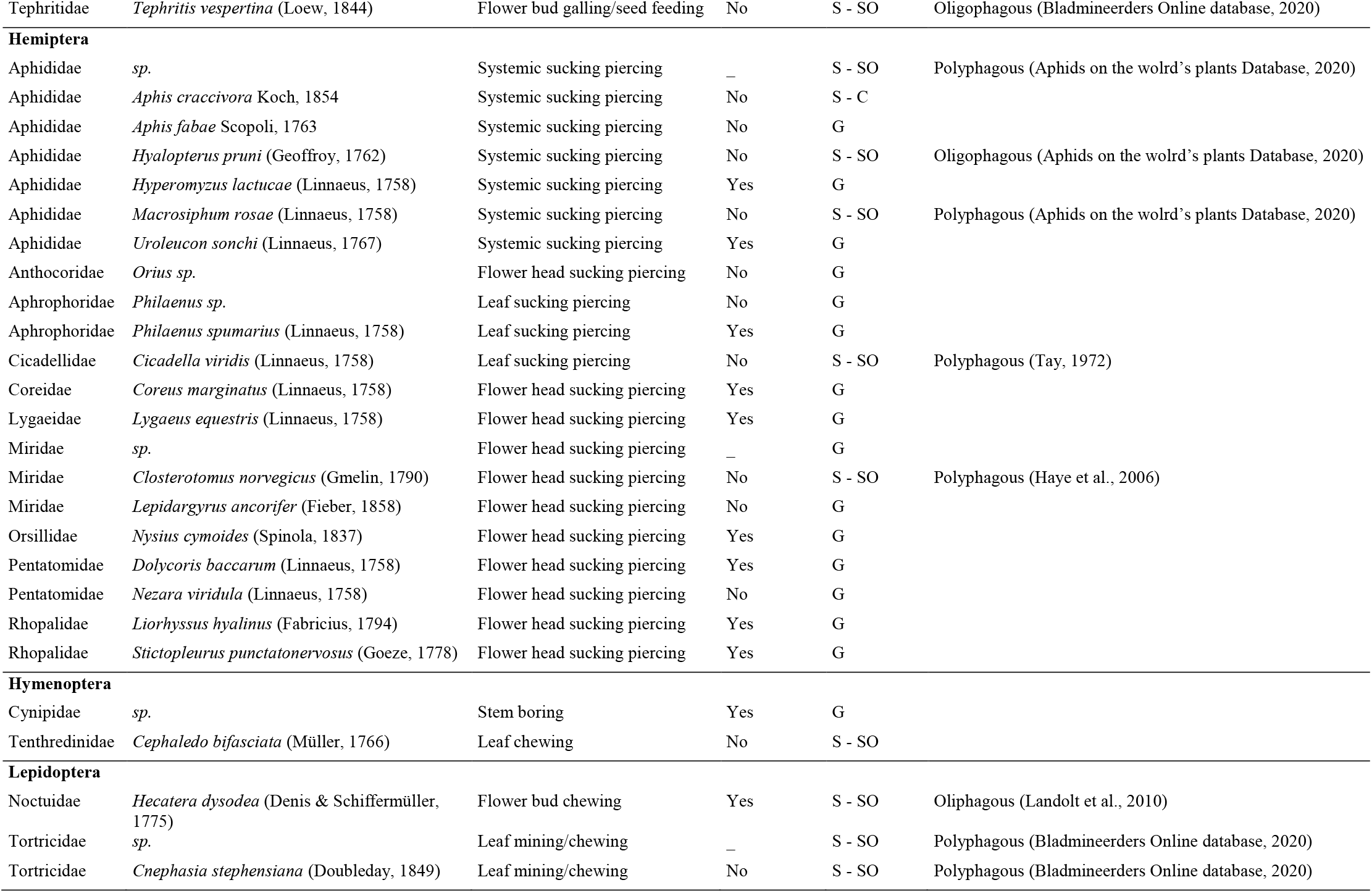
Herbivores from *Sonchus oleraceus* (SO) collected and identified through intense field sampling in three bioclimatic regions (Semi-oceanic, Mediterranean and Continental) in France in spring 2018. The field host range of these herbivores was defined by network analysis (S - SO: Specific to SO, S – S: Specific to the genus Sonchus, S – C: Specific to the tribe Cichorieae, G: Generalist, ?: Unknown host range). The column 3-years surveys refers to the species collected following classical procedures of biocontrol program (see Materials and Methods section).

The analysis of the subnetwork (Figure 2) also indicated that the herbivores collected on *S. oleraceus* were a resource for diverse natural enemies. In particular, 19 of the 47 herbivorous taxa collected were attacked by several species of parasitoid (12 species from the family Braconidae, 1 from Figitidae, and 1 from Ichneumonidae) and predators (6 Arachnida species, 1 from Cantharidae, 2 from Coccinnellidae, 3 from Syrphidae and 1 from Orthoptera). Moreover, among the 17 arthropods identified as candidate BCA for their restricted ecological host range, we detected interactions with natural enemies for six of them, one exhibiting interactions with 8 taxa from higher trophic levels (Figure 3).

Eventually, molecular analyses revealed particular patterns of omnivory involving several species from Heteroptera. We distinguished between intermediate omnivores (species feeding on both plants and herbivores, such as members of the Tephritidae and Aphididae), and top omnivores (species feeding on herbivores and natural enemies, such as members of the Syrphidae). The list of the taxa included in the subnetwork and of all the trophic interactions (*i.e*. the edge list) used to generate Figure 3 are provided in Supplementary Tables 3 and 4, respectively.

#### Assessing the contribution of the approach for the biocontrol program

The analysis of trophic interactions identified 47 taxa feeding on *S. oleraceus*, 37 of which were identified to species level. Nineteen of these 37 species had already been sampled in classical field surveys, the other 18 species being newly reported as herbivores of *S. oleraceus* in this CBC program (Table 2).

## Discussion

### The combination of observation and molecular data performs well for the characterization of interactions

The metabarcoding approach used made it possible to target a broad diversity of taxa (90 arthropod families, with identification to species level of 84% of the variants), as expected with the use of multiple markers (Alberdi et al., 2018; Creedy et al., 2019; Marquina et al., 2019). The combination of taxonomic assignments with subsequent observational data and information available in the literature was essential: 1) to validate the trophic links (predation and parasitism) and 2) to complement the identification in cases of failed amplification or taxonomic assignment, as advocated in other contexts (Derocles et al., 2018; Wirta et al., 2014). In the meta-network, 60% of the taxa interacted, suggesting that our methods performed very well for the reconstruction of interactions. More specifically, interaction detection rates obtained for plant-herbivores and herbivores-natural showed higher values than those usually reported in comparable contexts (Braukmann et al., 2017; Clare, 2014; Erickson et al., 2017; García-Robledo et al., 2013; Roslin and Majaneva, 2016). The high rate of interaction reported here for herbivorous arthropods can be explained by our decision to focus on intensive plant dissection and morphological determination. Retaining feeding interactions only after verification reduced the risk of false positives, over-estimating species interactions, related to the use of co-occurrence data (*i.e*. tourist insects on plants rather than actual trophic links) (Zhu et al., 2019). For arthropods, the rather low rate of natural enemies positive for preys can be multifactorial; *e.g*. mismatch between the primer pairs used and the prey species, low sequencing depth given the DNA yield ratio between consumer and prey, and degradation of DNA from consumed preys (Hosseini et al., 2008; Macías-Hernández et al., 2018; Sheppard and Harwood, 2005; Sheppard et al., 2004).

### A complementary inventory of herbivores feeding on *S. oleraceus*

The 47 taxa collected from *S. oleraceus*, covered a wide range of trophic guilds. This evidenced that the method herein employed did not bias the selection towards a particular trophic guild but allows the detection of herbivores exhibited diversified feeding habits. The selection of one or several BCA from these trophic guilds could offer a good complementarity of actions (Buccellato et al., 2019). Although the guild of flower head sucker-piercer was the richest, further dedicated experiments would be necessary to assess whether those candidate agents actually provide the best regulation action of *S. oleraceus* (Morin et al., 2009).

Several elements indicated that our approach is a good complement of the classical survey. For example, we provided a more detailed description of pollen chewers (only *Brassicogethes aeneus* (Fabricius, 1775) was recorded in the classical survey). For Diptera, the Tephritidae flies, *Tephritis cometa* Loew and *T. vespertina* Loew, are newly recorded. A phylogeny of this taxa based on the CO1 barcode showed that those species are closely related to *T. formosa* Loew (Smit et al., 2013), and we cannot, therefore, rule out possible molecular misidentification due to the short CO1 barcode used or host race differentiation, as frequently observed in this group (Diegisser et al., 2006). During the classical survey, despite intensive collections, the Tephritidae species *Campiglossa producta* has been sampled only on *S. oleraceus* from the Canary Islands. The intensive sampling efforts put in the present study on *S. oleraceus* led to the collection of rare *C. producta* specimens (only 15 specimens from two sites) in the continental bioclimatic region in France. This highlights that, despite a reduced area prospected and a limited time frame, some rare species were sampled and their ecological host range described. We acknowledge we missed some species occurring later in the season or out of the sampled area. For example, *Cystiphora sonchi* Vallot (Diptera: Cecidomyiidae) was collected in the classical surveys and passed specificity tests (Lesieur et al., 2020), but was not sampled from *S. oleraceus* in the three bioclimatic regions from April to June. This lack of detection in our sampling campaign was expected, as rates of infestation with this species peak in summer (Rizzo and Massa, 1998). Hence, this study should be regarded as a complement of usual procedures. However, with the rapid development of molecular technologies and associated drop in price (Kennedy et al., 2020), we believe that this approach will be soon applicable at larger sampling scales.

### Selection of BCA

#### First criterion: a restricted ecological host range

Based on the present results, 15 of the 47 herbivores feeding on *S. oleraceus* seemed to have an ecological host range restricted to *S. oleraceus*, and another two taxa appeared to be restricted to the genus *Sonchus*. All these taxa recovered from the ecological network are, thus, of particular interest as candidate BCA. However, contradictions were observed between the ecological host range described by network analysis and published findings or specificity test results (Lesieur et al., in prep). These discrepancies may be due to insufficient sampling for the recording of species interactions (as shown by accumulation curve on total interactions) or to the presence of cryptic host races or cryptic species that have yet to be deciphered. Additional studies will be required to characterize the ecological host ranges of these species further. In particular, two species, *Liriomyza sonchi* Hendel and *Ensina sonchi L*., were found associated with *S. oleraceus* and *S. asper* and were, therefore, considered to be candidate BCA because these plants are both invasive weeds in Australia (Cullen et al., 2012). However, a wider range of food resources has been reported in literature for these two species (Table 2). Conversely, the promising galling insect *T. formosa* passed specificity tests during the traditional phase of the CBC program. It was found to be restricted to the genus *Sonchus*, contrary to the results reported here, as we found *T. formosa* on *Crepis vesicaria* L. (belonging to Cripidinae, the same tribe but a different subtribe to *S. oleraceus*.). This plant was not indented to be tested as a potential food plant for *T. formosa*, and these results therefore highlight the complementarity of the ecological network approach for clarifying herbivore host range.

Moreover, in the interaction network, *C. producta* was identified on three different plant species from the Chicorieae tribe. This species, found only on *S. oleraceus* in the Canary Islands during the classical surveys, was considered a promising BCA for testing. The results presented here indicate that its host range would not be compatible with its use as a BCA, potentially leading to its exclusion from the list of candidate BCA. This example shows how the network developed here is complementary to classical procedures, making it possible to narrow down the list of candidate BCA to be tested.

The same applies to *Cheilosia latifrons* Zetterstedt, a species collected in the classical survey. In our study, we did not sample this species on *S. oleraceus*, but the meta-network indicated it was collected from *S. asper* and *Picris echioides* (L.), revealing its oligophagous dietary behavior. However, little is known about the biology and the host plants of *C. latifrons* (Schmid and Grossmann, 1996) and its taxonomy seems to be unsolved, calling into question the existence of a species complex defined on the basis of host plant use (Speight, 2014). We also observed discrepancies for specimens from Cynipidae that appeared to be generalist herbivores in the interaction network (associated with both *S. oleraceus* and *Carduus pycnocephalus* Spreng.), whereas subsequent analysis of the variants assigned to Cynipidae indicated a genetic structure more consistent with multiple cryptic species potentially specializing on the host plants from which they were collected. One species from Cynipidae is a known stem galler of *S. oleraceus: Aulacidea follioti* Barbotin (Bladmineerders Online database, 2020). However, this species is not yet present in any of the barcoding databases used here and could therefore only be assigned to family level. Further prospections to collect other Cynipidae specimens and rear them to adulthood would be required to confirm this identification.

#### Second criterion: limited interactions with natural enemies

By using metabarcoding to reconstruct interactions between arthropods, we were able to detect a wide range of parasitoids from their herbivore hosts, and some predators. We detected omnivorous dietary behavior in several groups from Heteroptera. Opportunistic predation through carnivory is common in Lygaeidae (Burdfield-Steel and Shuker, 2014) to supplement the low levels of protein supplied by plants. Carnivory has also been reported in Miridae (Wheeler, 2001) and sometimes leads to intraguild predation interactions. We found that both Syrphidae (Diptera) and Miridae (Heteroptera) fed on aphid species, but we also revealed that mirids could prey upon syrphids, as already demonstrated in arena experiments (Fréchette et al., 2006). Members of the Lygaeidae and Miridae were also found to be able to access and feed on larval stages of several Tephritidae species whilst inside the flower heads of *S. oleraceus*. This interaction does not seem to have been observed before and provides insight useful not only for the CBC program against *S. oleraceus*, but also with direct implications for other biological control programs, particularly those involving the conservation biological control of insect pests.

More specifically, among candidate BCA exhibiting a restricted ecological host range, some were associated to an important diversity of natural enemies, and should be considered of lower priority for testing (*i.e. Ensina sonchĩ*). We suggest that priority should be given to agent predated by a minimum parasitoid and predator species, and, when they exist, to an agent that possesses enemies from the most distant taxonomical group from those occurring in the range of introduction (Ollivier et al., 2020). It has been shown that newly created interactions between hosts and parasitoids in the introduced range are predictable based on the realised interactions in native range (Paynter et al., 2017; Veldtman et al., 2011). Further steps in this program would consist in investigating the diversity of natural enemies occurring in the range of invasion to anticipate new potential interactions and refine BCA choice.

## Conclusion

We demonstrate here the potential of network ecology for characterizing candidate BCA and their ecological interactions in the field. This characterization clearly benefited from the use of complementary approaches (morphological and molecular analyses) to identify plant/arthropod and arthropod/arthropod interactions and provided a solid framework for the establishment of an inventory of herbivores feeding on the target weed, their realized host range and interactions with natural enemies. Avenues for further investigation have been identified and in-depth studies are now required. The strength of this approach also lies in its capacity to screen field host ranges for multiple herbivore species simultaneously, without the need for as many tests as species. This potential to narrow down the list of candidate BCA for testing should help to save both time and money. Finally, in addition to the potential value of ecological network analysis to the CBC targeting Common Sowthistle in Australia, the data reported here are potentially useful for other future programs.

## Authors’ contributions

Conceptualization: M. Ollivier, V. Lesieur, M. S. Tixier, J.-F. Martin

Methodology: M. Ollivier, J. Tavoillot, F. Bénetière, V. Lesieur, J.-F. Martin

Data acquisition: M. Ollivier, J. Tavoillot, F. Bénetière, V. Lesieur, J.-F. Martin

Data analysis: M. Ollivier

Supervision: M. S. Tixier, J.-F. Martin,

Writing original draft: M. Ollivier

Writing review and editing: M. Ollivier, J. Tavoillot, V. Lesieur, M. S. Tixier, J.-F. Martin

## Acknowledgments

We are particularly grateful to A Coeur d’acier, G Delvare, B Derepas, J Harran, B Michel, A Migeon, E Pierre, JM Ramel and JC Streito for providing their taxonomic expertise for arthropod identification for construction of the local database. We also thank G Fried for assistance with the identification of some plant specimens. We also warmly thank all our coworkers involved in field sampling: P Audiot, M Corbin, R Guilhot, A Loiseau, L Olazcuaga, T Thomann, and N Vieira. This project is supported by funding from the Australian Government Department of Agriculture and Water Resources, as part of its Rural R&D for Profit programme, through AgriFutures Australia (Rural Industries Research and Development Corporation) (PRJ--010527).

