## Supplementary materials for "Metabarcoding and ecological interaction networks for selecting candidate biological control agents"

### Supplementary Texts

### Supplementary Tables

### Supplementary Figures

References

### Supplementary Texts

*Text 1: Curation and transformation of the interaction matrix*

In order to produce an interaction matrix, in which observations are resources and variables are consumers, the original dataset should be checked. First, for each sample, the taxa having the highest number of reads were set as the consumer. This assumption was confirmed in most of the cases (the specimen collected in the tube was the consumer and produced the highest number of reads). However due to the simultaneous detection of possible preys and parasitoids from the specimen sampled and the contrasted amplification rates between taxonomic groups (Alberdi et al. 2018), this was not always the case. Consequently, a second step of data manual validation was required on each sample (Supplementary Figure S2) in order to 1) verify whether the consumer was correctly defined according to the observation from the field, 2) add the source plant for herbivorous arthropods, 3) define the status of the taxa simultaneously amplified with the consumer (prey, parasitoid or contamination). The next step consisted in removing cannibalistic interactions (i. e. delete reads at the intersection of a same source and consumer species). Indeed, DNA-based methods are not suitable to disentangle predator and prey sequences (Taberlet et al. 2018). As metabarcoding neither provides quantitative information about the trophic links (Deagle et al. 2019), the numbers of reads were converted to binary information, then occurrence frequencies of interactions between a pair of consumer/prey species were calculated as the percentage of consumer samples positive for the prey over the total number of consumer sampled in the dataset. All analyses were carried out under R version 3.5.2 (R Core Team 2018).

### Supplementary Tables

Table 1: Locations of the different sites prospected per sampling session in the three bioclimatic regions.

|  | **Site** | **Latitude** | **Longitude** | **Environment** |
| --- | --- | --- | --- | --- |
| **Semi-oceanic** |  |  |  |  |
| *Session 1* | Laurabuc | 43.26462 | 1.99578 | Agricultural |
|  | Saint-Martin-Lalande | 43.29428 | 1.99238 | Agricultural |
|  | Castelnaudary | 43.33395 | 1.99262 | Agricultural |
| *Session 2* | Laurabuc | 43.26462 | 1.99578 | Agricultural |
|  | Saint-Martin-Lalande | 43.29428 | 1.99238 | Agricultural |
| *Session 3* | Laurabuc | 43.26462 | 1.99578 | Agricultural |
|  | Villasavary | 43.23643 | 2.03641 | Ruderal |
| **Mediterranean** |  |  |  |  |
| *Session 1* | La Valsière | 43.65123 | 3.83356 | Ruderal |
|  | La Valsière | 43.65123 | 3.83356 | Ruderal |
| *Session 2* | Mauguio | 43.6069 | 3.98868 | Agricultural |
|  | La Valsière | 43.65123 | 3.83356 | Ruderal |
| *Session 3* | Mauguio | 43.6069 | 3.98868 | Agricultural |
|  | La Valsière | 43.65123 | 3.83356 | Ruderal |
| **Continental** |  |  |  |  |
| *Session 1* | Mornant | 45.60496 | 4.68696 | Agricultural |
|  | Saint-Génis-Laval | 45.6909 | 4.76503 | Agricultural |
| *Session 2* | Saint Laurent d’Agny | 45.6335 | 4.6831 | Agricultural |
|  | Le Clair | 45.63451 | 4.69026 | Ruderal |
| *Session 3* | Saint Laurent d’Agny | 45.6335 | 4.6831 | Agricultural |
|  | Thurigneux | 45.59188 | 4.64755 | Ruderal |

| **Marker** | **Raw reads** | **Filtered reads** | **Percentage of reads retained** | **Variants 2%** |
| --- | --- | --- | --- | --- |
| **HEX** | 9083407 | 1205146 | 0,133 | 408 |
| **HCO** | 48226250 | 7190702 | 0,149 | 847 |
| **LEP** | 50173753 | 12320157 | 0,246 | 759 |
| **Total** | **107483410** | **20716005** | **0,193** | **2014** |

Table 2: Detailed number of reads obtained from high-throughput sequencing for each of
the CO1 markers employed

Table 3: List of the taxa included in the sub-network (Figure 2)

| **Node ID** | ***Taxon*** | **Order (Family)** | **Trophic level** |
| --- | --- | --- | --- |
| **1** | *Agapanthia cardui* | Coleoptera (Cerambycidae) | Herbivore |
| **2** | *Agapanthia pannonica* | Coleoptera (Cerambycidae) | Herbivore |
| **3** | *Agromyzidae* | Diptera (Agromyzidae) | Herbivore |
| **4** | *Alloxysta consobrina* | Hymeoptera (Figitidae) | Natural enemy (parasitoid) |
| **5** | *Anthemis altissima* | Asterales (Asteraceae) | Plant |
| **6** | *Aphididae* | Hemiptera (Aphididae) | Herbivore |
| **7** | *Aphidius ervi* | Hemiptera (Aphididae) | Herbivore |
| **8** | *Aphidius sonchi* | Hemiptera (Aphididae) | Herbivore |
| **9** | *Aphis craccivora* | Hemiptera (Aphididae) | Herbivore |
| **10** | *Aphis fabae* | Hemiptera (Aphididae) | Herbivore |
| **11** | *Avena sp.* | Poales (Poaceae) | Plant |
| **12** | *Bracon radialis* | Hymeoptera (Braconidae) | Natural enemy (parasitoid) |
| **13** | *Bracon sp.* | Hymeoptera (Braconidae) | Natural enemy (parasitoid) |
| **14** | *Bracon telengai* | Hymeoptera (Braconidae) | Natural enemy (parasitoid) |
| **15** | *Braconidae* | Hymeoptera (Braconidae) | Natural enemy (parasitoid) |
| **16** | *Brassicogethes aeneus* | Coleoptera (Nitidulidae) | Herbivore |
| **17** | *Campiglossa producta* | Diptera (Tephritidae) | Herbivore |
| **18** | *Carduus pycnocephalus* | Asterales (Asteraceae) | Plant |
| **19** | *Cerastium glomeratum* | Caryophyllales (Caryophyllaceae) | Plant |
| **20** | *Cicadella viridis* | Hemiptera (Cicadellidae) | Herbivore |
| **21** | *Cichorium intybus* | Asterales (Asteraceae) | Plant |
| **22** | *Cirsium arvense* | Asterales (Asteraceae) | Plant |
| **23** | *Cirsium vulgare* | Asterales (Asteraceae) | Plant |
| **24** | *Closterotomus norvegicus* | Hemiptera (Miridae) | Herbivore |
| **25** | *Cnephasia stephensiana* | Lepidoptera (Tortricidae) | Herbivore |
| **26** | *Coccinella septempunctata* | Coleoptera (Coccinellidae) | Natural enemy (predator) |
| **27** | *Coreus marginatus* | Hemiptera (Coreidae) | Herbivore |
| **28** | *Crepis foetida* | Asterales (Asteraceae) | Plant |
| **29** | *Crepis vesicaria* | Asterales (Asteraceae) | Plant |
| **30** | *Curculionidae* | Coleoptera (Curculionidae) | Herbivore |
| **31** | *Cynipidae* | Hymenoptera (Cynipidae) | Herbivore |
| **32** | *Dacnusa maculipes* | Hymenoptera (Braconidae) | Natural enemy (parasitoid) |
| **33** | *Dasytes tristiculus* | Coleoptera (Melyridae) | Herbivore |
| **34** | *Diadegma erucator* | Hymenoptera (Ichneumonidae) | Natural enemy (parasitoid) |
| **35** | *Dicot unknown* | / | Plant |
| **36** | *Diplazon tibiatorius* | Hymenoptera (Ichneumonidae) | Natural enemy (parasitoid) |
| **37** | *Dolycoris baccarum* | Hemiptera (Pentatomidae) | Herbivore |
| **38** | *Ensina sonchi* | Diptera (Tephritidae) | Herbivore |
| **39** | *Ephedrus sp.* | Hymenoptera (Braconidae) | Natural enemy (parasitoid) |
| **40** | *Episyrphus balteatus* | Diptera (Syrphidae) | Natural enemy (predator) |
| **41** | *Erigeron sp.* | Asterales (Asteraceae) | Plant |
| **42** | *Euphorbia helioscopia* | Malpighiales (Euphorbiceae) | Plant |
| **43** | *Geranium rotundifolium* | Geraniales (Geraniaceae) | Plant |
| **44** | *Geranium sp.* | Geraniales (Geraniaceae) | Plant |
| **45** | *Hecatera dysodea* | Lepidoptera (Noctuidae) | Herbivore |
| **46** | *Heliophanus cupreus* | Araneae (Saltividae) | Natural enemy (predator) |
| **47** | *Heliophanus sp.* | Araneae (Saltividae) | Natural enemy (predator) |
| **48** | *Hordeum murinum* | Poales (Poaceae) | Plant |
| **49** | *Hyalopterus pruni* | Hemiptera (Aphididae) | Herbivore |
| **50** | *Hypericum perforatum* | Theales (Clusiaceae) | Plant |
| **51** | *Hyperomyzus lactucae* | Hemiptera (Aphididae) | Herbivore |
| **52** | *Hypochaeris radicata* | Asterales (Asteraceae) | Plant |
| **53** | *Knautia arvensis* | Dipsacales (Dipsacaceae) | Plant |
| **54** | *Kochiura aulica* | Araneae (Theridiidae) | Natural enemy (predator) |
| **55** | *Lactuca serriola* | Asterales (Asteraceae) | Plant |
| **56** | *Lathyrus sp.* | Fabales (Fabaceae) | Plant |
| **57** | *Lepidargyrus ancorifer* | Hemiptera (Miridae) | Herbivore |
| **58** | *Lepidium draba* | Capparales (Brassicaceae) | Plant |
| **59** | *Leptophyes punctatissima* | Orthoptera (Tettigoniidae) | Natural enemy (predator) |
| **60** | *Liorhyssus hyalinus* | Hemiptera (Rhopalidae) | Omnivore |
| **61** | *Lipolexis gracilis* | Hymenoptera (Braconidae) | Natural enemy (parasitoid) |
| **62** | *Liriomyza sonchi* | Diptera (Agromyzidae) | Herbivore |
| **63** | *Lixus punctiventris* | Coleoptera (Curculionidae) | Herbivore |
| **64** | *Lolium sp.* | Poales (Poaceae) | Plant |
| **65** | *Lygaeus equestris* | Hemiptera (Lygaeidae) | Omnivore |
| **66** | *Macrosiphum rosae* | Hemiptera (Aphididae) | Herbivore |
| **67** | *Medicago orbicularis* | Fabales (Fabaceae) | Plant |
| **68** | *Microplitis sp.* | Hymenoptera (Braconidae) | Natural enemy (parasitoid) |
| **69** | *Miridae* | Hemiptera (Miridae) | Omnivore |
| **70** | *Misumena vatia* | Araneae (Thomisidae) | Omnivore |
| **71** | *Neoscona adianta* | Araneae (Araneidae) | Omnivore |
| **72** | *Nezara viridula* | Hemiptera (Pentatomidae) | Herbivore |
| **73** | *Nysius cymoides* | Hemiptera (Orsillidae) | Omnivore |
| **74** | *Oedemera crassipes* | Coleoptera (Oedemeridae) | Herbivore |
| **75** | *Oedemera flavipes* | Coleoptera (Oedemeridae) | Herbivore |
| **76** | *Ophiomyia cunctata (beckeri)* | Diptera (Agromyzidae) | Herbivore |
| **77** | *Opiinae* | Hymenoptera (Braconidae) | Natural enemy (parasitoid) |
| **78** | *Orius sp.* | Hemiptera (Anthocoridae) | Herbivore |
| **79** | *Papaver rhoeas* | Ranunculales (Papaveraceae) | Plant |
| **80** | *Philaenus sp.* | Hemiptera (Aphrophoridae) | Herbivore |
| **81** | *Philaenus spumarius* | Hemiptera (Aphrophoridae) | Herbivore |
| **82** | *Phytomyza horticola (syngenesiae)* | Diptera (Agromyzidae) | Herbivore |
| **83** | *Phytomyza lateralis (cichorii)* | Diptera (Agromyzidae) | Herbivore |
| **84** | *Picris echioides* | Asterales (Asteraceae) | Plant |
| **85** | *Plantago lanceolata* | Lamiales (Plantaginaceae) | Plant |
| **86** | *Platycheirus scutatus* | Diptera (Syrphidae) | Natural enemy (predator) |
| **87** | *Poa pratensis* | Poales (Poaceae) | Plant |
| **88** | *Poa trivialis* | Poales (Poaceae) | Plant |
| **89** | *Poaceae* | Poales (Poaceae) | Plant |
| **90** | *Praon volucre* | Hymenoptera (Braconidae) | Natural enemy (parasitoid) |
| **91** | *Psilothrix viridicoeruleus* | Coleoptera (Dasytidae) | Herbivore |
| **92** | *Psyllobora vigintiduopunctata* | Coleoptera (Coccinellidae) | Herbivore |
| **93** | *Rhagonycha fulva* | Coleoptera (Cantharidae) | Natural enemy (predator) |
| **94** | *Rubus ulmifolius* | Rosales (Rosaceae) | Plant |
| **95** | *Rumex crispus* | Polygonales (Polygonaceae) | Plant |
| **96** | *Runcinia grammica* | Araneae (Thomisidae) | Omnivore |
| **97** | *Senecio vulgaris* | Asterales (Asteraceae) | Plant |
| **98** | *Sonchus asper* | Asterales (Asteraceae) | Plant |
| **99** | *Sonchus oleraceus* | Asterales (Asteraceae) | Plant |
| **100** | *Sphaerophoria sp.* | Diptera (Syrphidae) | Plant |
| **101** | *Stictopleurus punctatonervosus* | Hemiptera (Rhopalidae) | Omnivore |
| **102** | *Taraxacum officinale* | Asterales (Asteraceae) | Plant |
| **103** | *Cephaledo bifasciata* | Hymenoptera (Tenthredinidae) | Herbivore |
| **104** | *Tephritidae* | Diptera (Tephritidae) | Herbivore |
| **105** | *Tephritis cometa* | Diptera (Tephritidae) | Herbivore |
| **106** | *Tephritis formosa* | Diptera (Tephritidae) | Herbivore |
| **107** | *Tephritis sp.* | Diptera (Tephritidae) | Herbivore |
| **108** | *Tephritis vespertina* | Diptera (Tephritidae) | Herbivore |
| **109** | *Tortricidae* | Lepidoptera (Tortricidae) | Herbivore |
| **110** | *Trombidiidae* | Acariformes (Trombidiidae) | Natural enemy (Parasit) |
| **111** | *Tytthaspis sedecimpunctata* | Coleoptera (Coccinellidae) | Omnivore |
| **112** | *Uroleucon sonchi* | Hemiptera (Aphididae) | Herbivore |
| **113** | *Verbascum blattaria* | Scrophulariales (Scrophulariaceae) | Plant |
| **114** | *Veronica persica* | Lamiales (Plantaginaceae) | Plant |
| **115** | *Vicia sativa* | Fabales (Fabaceae) | Plant |
| **116** | *Viola arvensis* | Violales (Violaceae) | Plant |

Table 4: List of interactions included in the sub-network (Figure 2)

| **Interaction** | **Resource** | **Consumer** | **Occurrence frequency** |
| --- | --- | --- | --- |
| **1** | *Sonchus asper* | *Agromyzidae* | 20 |
| **2** | *Sonchus asper* | *Campiglossa producta* | 13 |
| **3** | *Sonchus asper* | *Ensina sonchi* | 9 |
| **4** | *Sonchus asper* | *Hyperomyzus lactucae* | 17 |
| **5** | *Sonchus asper* | *Liorhyssus hyalinus* | 34 |
| **6** | *Sonchus asper* | *Liriomyza sonchi* | 14 |
| **7** | *Sonchus asper* | *Lixus punctiventris* | 2 |
| **8** | *Sonchus asper* | *Ophiomyia cunctata (beckeri)* | 8 |
| **9** | *Sonchus asper* | *Philaenus spumarius* | 9 |
| **10** | *Sonchus asper* | *Phytomyza horticola (syngenesiae)* | 15 |
| **11** | *Sonchus asper* | *Psilothrix viridicoeruleus* | 9 |
| **12** | *Sonchus asper* | *Tephritis formosa* | 7 |
| **13** | *Sonchus asper* | *Tephritis sp* | 11 |
| **14** | *Sonchus oleraceus* | *Agapanthia cardui* | 15 |
| **15** | *Sonchus oleraceus* | *Agapanthia pannonica* | 50 |
| **16** | *Sonchus oleraceus* | *Agromyzidae* | 52 |
| **17** | *Sonchus oleraceus* | *Aphididae* | 100 |
| **18** | *Sonchus oleraceus* | *Aphis craccivora* | 50 |
| **19** | *Sonchus oleraceus* | *Aphis fabae* | 71 |
| **20** | *Sonchus oleraceus* | *Brassicogethes aeneus* | 21 |
| **21** | *Sonchus oleraceus* | *Campiglossa producta* | 60 |
| **22** | *Sonchus oleraceus* | *Cicadella viridis* | 100 |
| **23** | *Sonchus oleraceus* | *Closterotomus norvegicus* | 100 |
| **24** | *Sonchus oleraceus* | *Cnephasia stephensiana* | 33 |
| **25** | *Sonchus oleraceus* | *Curculionidae* | 12 |
| **26** | *Sonchus oleraceus* | *Cynipidae* | 50 |
| **27** | *Sonchus oleraceus* | *Dasytes tristiculus* | 8 |
| **28** | *Sonchus oleraceus* | *Dolycoris baccarum* | 50 |
| **29** | *Sonchus oleraceus* | *Ensina sonchi* | 90 |
| **30** | *Sonchus oleraceus* | *Hecatera dysodea* | 100 |
| **31** | *Sonchus oleraceus* | *Hyalopterus pruni* | 100 |
| **32** | *Sonchus oleraceus* | *Hyperomyzus lactucae* | 69 |
| **33** | *Sonchus oleraceus* | *Lepidargyrus ancorifer* | 22 |
| **34** | *Sonchus oleraceus* | *Liorhyssus hyalinus* | 50 |
| **35** | *Sonchus oleraceus* | *Liriomyza sonchi* | 79 |
| **36** | *Sonchus oleraceus* | *Lixus punctiventris* | 96 |
| **37** | *Sonchus oleraceus* | *Lygaeus equestris* | 11 |
| **38** | *Sonchus oleraceus* | *Macrosiphum rosae* | 100 |
| **39** | *Sonchus oleraceus* | *Miridae* | 38 |
| **40** | *Sonchus oleraceus* | *Nezara viridula* | 33 |
| **41** | *Sonchus oleraceus* | *Nysius cymoides* | 12 |
| **42** | *Sonchus oleraceus* | *Oedemera crassipes* | 100 |
| **43** | *Sonchus oleraceus* | *Oedemera flavipes* | 33 |
| **44** | *Sonchus oleraceus* | *Ophiomyia cunctata (beckeri)* | 88 |
| **45** | *Sonchus oleraceus* | *Orius sp* | 50 |
| **46** | *Sonchus oleraceus* | *Philaenus sp* | 25 |
| **47** | *Sonchus oleraceus* | *Philaenus spumarius* | 40 |
| **48** | *Sonchus oleraceus* | *Phytomyza horticola (syngenesiae)* | 63 |
| **49** | *Sonchus oleraceus* | *Phytomyza lateralis (cichorii)* | 50 |
| **50** | *Sonchus oleraceus* | *Psilothrix viridicoeruleus* | 18 |
| **51** | *Sonchus oleraceus* | *Stictopleurus punctatonervosus* | 57 |
| **52** | *Sonchus oleraceus* | *Cephaledo bifasciata* | 100 |
| **53** | *Sonchus oleraceus* | *Tephritidae* | 100 |
| **54** | *Sonchus oleraceus* | *Tephritis cometa* | 100 |
| **55** | *Sonchus oleraceus* | *Tephritis formosa* | 89 |
| **56** | *Sonchus oleraceus* | *Tephritis sp* | 48 |
| **57** | *Sonchus oleraceus* | *Tephritis vespertina* | 100 |
| **58** | *Sonchus oleraceus* | *Tortricidae* | 100 |
| **59** | *Sonchus oleraceus* | *Uroleucon sonchi* | 84 |
| **60** | *Sonchus oleraceus* | *Coreus marginatus* | 60 |
| **61** | *Poa pratensis* | *Philaenus spumarius* | 7 |
| **62** | *Cirsium arvense* | *Aphis fabae* | 14 |
| **63** | *Cirsium arvense* | *Brassicogethes aeneus* | 54 |
| **64** | *Cirsium arvense* | *Oedemera flavipes* | 33 |
| **65** | *Cirsium arvense* | *Phytomyza horticola (syngenesiae)* | 2 |
| **66** | *Poaceae* | *Philaenus spumarius* | 1 |
| **67** | *Poaceae* | *Uroleucon sonchi* | 1 |
| **68** | *Picris echioides* | *Agromyzidae* | 1 |
| **69** | *Picris echioides* | *Brassicogethes aeneus* | 7 |
| **70** | *Picris echioides* | *Hyperomyzus lactucae* | 4 |
| **71** | *Picris echioides* | *Lepidargyrus ancorifer* | 11 |
| **72** | *Picris echioides* | *Liorhyssus hyalinus* | 3 |
| **73** | *Picris echioides* | *Nezara viridula* | 17 |
| **74** | *Picris echioides* | *Philaenus spumarius* | 9 |
| **75** | *Picris echioides* | *Phytomyza horticola (syngenesiae)* | 15 |
| **76** | *Picris echioides* | *Phytomyza lateralis (cichorii)* | 33 |
| **77** | *Picris echioides* | *Tephritis sp* | 7 |
| **78** | *Picris echioides* | *Uroleucon sonchi* | 1 |
| **79** | *Lolium sp* | *Coreus marginatus* | 20 |
| **80** | *Veronica persica* | *Philaenus sp* | 25 |
| **81** | *Euphorbia helioscopia* | *Philaenus spumarius* | 1 |
| **82** | *Dicot unknown* | *Curculionidae* | 12 |
| **83** | *Dicot unknown* | *Philaenus sp* | 25 |
| **84** | *Lactuca serriola* | *Agromyzidae* | 23 |
| **85** | *Lactuca serriola* | *Aphis craccivora* | 50 |
| **86** | *Lactuca serriola* | *Hyperomyzus lactucae* | 6 |
| **87** | *Lactuca serriola* | *Liorhyssus hyalinus* | 3 |
| **88** | *Lactuca serriola* | *Lixus punctiventris* | 2 |
| **89** | *Lactuca serriola* | *Ophiomyia cunctata (beckeri)* | 4 |
| **90** | *Lactuca serriola* | *Philaenus spumarius* | 1 |
| **91** | *Lactuca serriola* | *Phytomyza horticola (syngenesiae)* | 4 |
| **92** | *Lactuca serriola* | *Phytomyza lateralis (cichorii)* | 17 |
| **93** | *Lactuca serriola* | *Uroleucon sonchi* | 5 |
| **94** | *Senecio vulgaris* | *Nezara viridula* | 17 |
| **95** | *Carduus pycnocephalus* | *Agapanthia cardui* | 62 |
| **96** | *Carduus pycnocephalus* | *Agromyzidae* | 1 |
| **97** | *Carduus pycnocephalus* | *Curculionidae* | 76 |
| **98** | *Carduus pycnocephalus* | *Cynipidae* | 50 |
| **99** | *Carduus pycnocephalus* | *Lepidargyrus ancorifer* | 44 |
| **100** | *Carduus pycnocephalus* | *Liorhyssus hyalinus* | 3 |
| **101** | *Carduus pycnocephalus* | *Miridae* | 29 |
| **102** | *Carduus pycnocephalus* | *Nezara viridula* | 17 |
| **103** | *Carduus pycnocephalus* | *Philaenus spumarius* | 6 |
| **104** | *Hordeum murinum* | *Philaenus spumarius* | 1 |
| **105** | *Cerastium glomeratum* | *Philaenus spumarius* | 1 |
| **106** | *Viola arvensis* | *Philaenus spumarius* | 1 |
| **107** | *Lathyrus sp* | *Philaenus spumarius* | 4 |
| **108** | *Geranium sp* | *Philaenus spumarius* | 3 |
| **109** | *Vicia sativa* | *Lygaeus equestris* | 11 |
| **110** | *Vicia sativa* | *Miridae* | 8 |
| **111** | *Vicia sativa* | *Philaenus spumarius* | 4 |
| **112** | *Anthemis altissima* | *Oedemera flavipes* | 33 |
| **113** | *Anthemis altissima* | *Psilothrix viridicoeruleus* | 9 |
| **114** | *Anthemis altissima* | *Tephritis sp* | 4 |
| **115** | *Avena sp* | *Miridae* | 4 |
| **116** | *Erigeron sp* | *Hyperomyzus lactucae* | 2 |
| **117** | *Erigeron sp* | *Psilothrix viridicoeruleus* | 18 |
| **118** | *Erigeron sp* | *Stictopleurus punctatonervosus* | 14 |
| **119** | *Papaver rhoeas* | *Brassicogethes aeneus* | 4 |
| **120** | *Papaver rhoeas* | *Miridae* | 4 |
| **121** | *Taraxacum officinale* | *Campiglossa producta* | 13 |
| **122** | *Taraxacum officinale* | *Phytomyza horticola (syngenesiae)* | 2 |
| **123** | *Taraxacum officinale* | *Stictopleurus punctatonervosus* | 14 |
| **124** | *Crepis vesicaria* | *Aphis fabae* | 14 |
| **125** | *Crepis vesicaria* | *Campiglossa producta* | 13 |
| **126** | *Crepis vesicaria* | *Tephritis formosa* | 3 |
| **127** | *Crepis vesicaria* | *Tephritis sp* | 26 |
| **128** | *Poa trivialis* | *Orius sp* | 50 |
| **129** | *Poa trivialis* | *Philaenus spumarius* | 1 |
| **130** | *Plantago lanceolata* | *Uroleucon sonchi* | 2 |
| **131** | *Cichorium intybus* | *Philaenus spumarius* | 1 |
| **132** | *Cichorium intybus* | *Uroleucon sonchi* | 1 |
| **133** | *Crepis foetida* | *Lepidargyrus ancorifer* | 11 |
| **134** | *Crepis foetida* | *Nysius cymoides* | 12 |
| **135** | *Knautia arvensis* | *Philaenus sp* | 25 |
| **136** | *Rumex crispus* | *Philaenus spumarius* | 3 |
| **137** | *Rubus ulmifolius* | *Philaenus spumarius* | 1 |
| **138** | *Geranium rotundifolium* | *Liorhyssus hyalinus* | 3 |
| **139** | *Geranium rotundifolium* | *Uroleucon sonchi* | 6 |
| **140** | *Hypericum perforatum* | *Coreus marginatus* | 20 |
| **141** | *Cirsium vulgare* | *Nysius cymoides* | 12 |
| **142** | *Lepidium draba* | *Nysius cymoides* | 38 |
| **143** | *Verbascum blattaria* | *Lepidargyrus ancorifer* | 11 |
| **144** | *Verbascum blattaria* | *Nysius cymoides* | 25 |
| **145** | *Medicago orbicularis* | *Dolycoris baccarum* | 17 |
| **146** | *Medicago orbicularis* | *Nezara viridula* | 17 |
| **147** | *Psilothrix viridicoeruleus* | *Neoscona adianta* | 33 |
| **148** | *Brassicogethes aeneus* | *Misumena vatia* | 17 |
| **149** | *Brassicogethes aeneus* | *Neoscona adianta* | 33 |
| **150** | *Brassicogethes aeneus* | *Runcinia grammica* | 23 |
| **151** | *Oedemera flavipes* | *Runcinia grammica* | 8 |
| **152** | *Ophiomyia cunctata (beckeri)* | *Braconidae* | 12 |
| **153** | *Phytomyza horticola (syngenesiae)* | *Bracon telengai* | 33 |
| **154** | *Phytomyza horticola (syngenesiae)* | *Dacnusa maculipes* | 100 |
| **155** | *Phytomyza horticola (syngenesiae)* | *Opiinae* | 100 |
| **156** | *Tephritidae* | *Bracon radialis* | 67 |
| **157** | *Tephritidae* | *Bracon sp* | 28 |
| **158** | *Tephritidae* | *Braconidae* | 12 |
| **159** | *Campiglossa producta* | *Bracon radialis* | 33 |
| **160** | *Campiglossa producta* | *Bracon sp* | 6 |
| **161** | *Campiglossa producta* | *Runcinia grammica* | 8 |
| **162** | *Ensina sonchi* | *Bracon sp* | 22 |
| **163** | *Ensina sonchi* | *Bracon telengai* | 33 |
| **164** | *Ensina sonchi* | *Braconidae* | 75 |
| **165** | *Ensina sonchi* | *Coreus marginatus* | 40 |
| **166** | *Ensina sonchi* | *Kochiura aulica* | 17 |
| **167** | *Ensina sonchi* | *Liorhyssus hyalinus* | 3 |
| **168** | *Ensina sonchi* | *Lygaeus equestris* | 56 |
| **169** | *Ensina sonchi* | *Nysius cymoides* | 12 |
| **170** | *Tephritis formosa* | *Bracon sp* | 44 |
| **171** | *Tephritis formosa* | *Bracon telengai* | 33 |
| **172** | *Tephritis formosa* | *Kochiura aulica* | 17 |
| **173** | *Tephritis formosa* | *Lygaeus equestris* | 11 |
| **174** | *Tephritis formosa* | *Miridae* | 4 |
| **175** | *Aphis fabae* | *Liorhyssus hyalinus* | 3 |
| **176** | *Aphis fabae* | *Lipolexis gracilis* | 100 |
| **177** | *Aphis fabae* | *Lygaeus equestris* | 11 |
| **178** | *Aphis fabae* | *Praon volucre* | 9 |
| **179** | *Hyalopterus pruni* | *Aphidius sonchi* | 4 |
| **180** | *Hyperomyzus lactucae* | *Alloxysta consobrina* | 50 |
| **181** | *Hyperomyzus lactucae* | *Aphidius sonchi* | 83 |
| **182** | *Hyperomyzus lactucae* | *Leptophyes punctatissima* | 20 |
| **183** | *Hyperomyzus lactucae* | *Miridae* | 8 |
| **184** | *Hyperomyzus lactucae* | *Praon volucre* | 86 |
| **185** | *Hyperomyzus lactucae* | *Sphaerophoria sp* | 11 |
| **186** | *Hyperomyzus lactucae* | *Stictopleurus punctatonervosus* | 14 |
| **187** | *Hyperomyzus lactucae* | *Trombidiidae* | 50 |
| **188** | *Hyperomyzus lactucae* | *Tytthaspis sedecimpunctata* | 14 |
| **189** | *Aphididae* | *Alloxysta consobrina* | 50 |
| **190** | *Aphididae* | *Platycheirus scutatus* | 100 |
| **191** | *Uroleucon sonchi* | *Aphidius ervi* | 100 |
| **192** | *Uroleucon sonchi* | *Coccinella septempunctata* | 14 |
| **193** | *Uroleucon sonchi* | *Ephedrus sp* | 80 |
| **194** | *Uroleucon sonchi* | *Episyrphus balteatus* | 6 |
| **195** | *Uroleucon sonchi* | *Heliophanus sp* | 50 |
| **196** | *Uroleucon sonchi* | *Praon volucre* | 5 |
| **197** | *Uroleucon sonchi* | *Psyllobora vigintiduopunctata* | 9 |
| **198** | *Uroleucon sonchi* | *Rhagonycha fulva* | 22 |
| **199** | *Philaenus spumarius* | *Heliophanus cupreus* | 33 |
| **200** | *Miridae* | *Ephedrus sp* | 10 |
| **201** | *Miridae* | *Runcinia grammica* | 8 |
| **202** | *Lepidargyrus ancorifer* | *Runcinia grammica* | 8 |
| **203** | *Liorhyssus hyalinus* | *Kochiura aulica* | 17 |
| **204** | *Hecatera dysodea* | *Microplitis sp* | 100 |
| **205** | *Tortricidae* | *Diadegma erucator* | 75 |
| **206** | *Hypochaeris radicata* | *Agromyzidae* | 1 |
| **207** | *Psyllobora vigintiduopunctata* | *Misumena vatia* | 17 |
| **208** | *Episyrphus balteatus* | *Misumena vatia* | 17 |
| **209** | *Platycheirus scutatus* | *Diplazon tibiatorius* | 100 |
| **210** | *Sphaerophoria sp* | *Lygaeus equestris* | 11 |
| **211** | *Sphaerophoria sp* | *Misumena vatia* | 50 |
| **212** | *Sphaerophoria sp* | *Neoscona adianta* | 67 |
| **213** | *Sphaerophoria sp* | *Runcinia grammica* | 15 |

### Supplementary Figures

Figure S1: Overview of the entire process for the reconstruction of interactions between plants, herbivores and natural enemies.

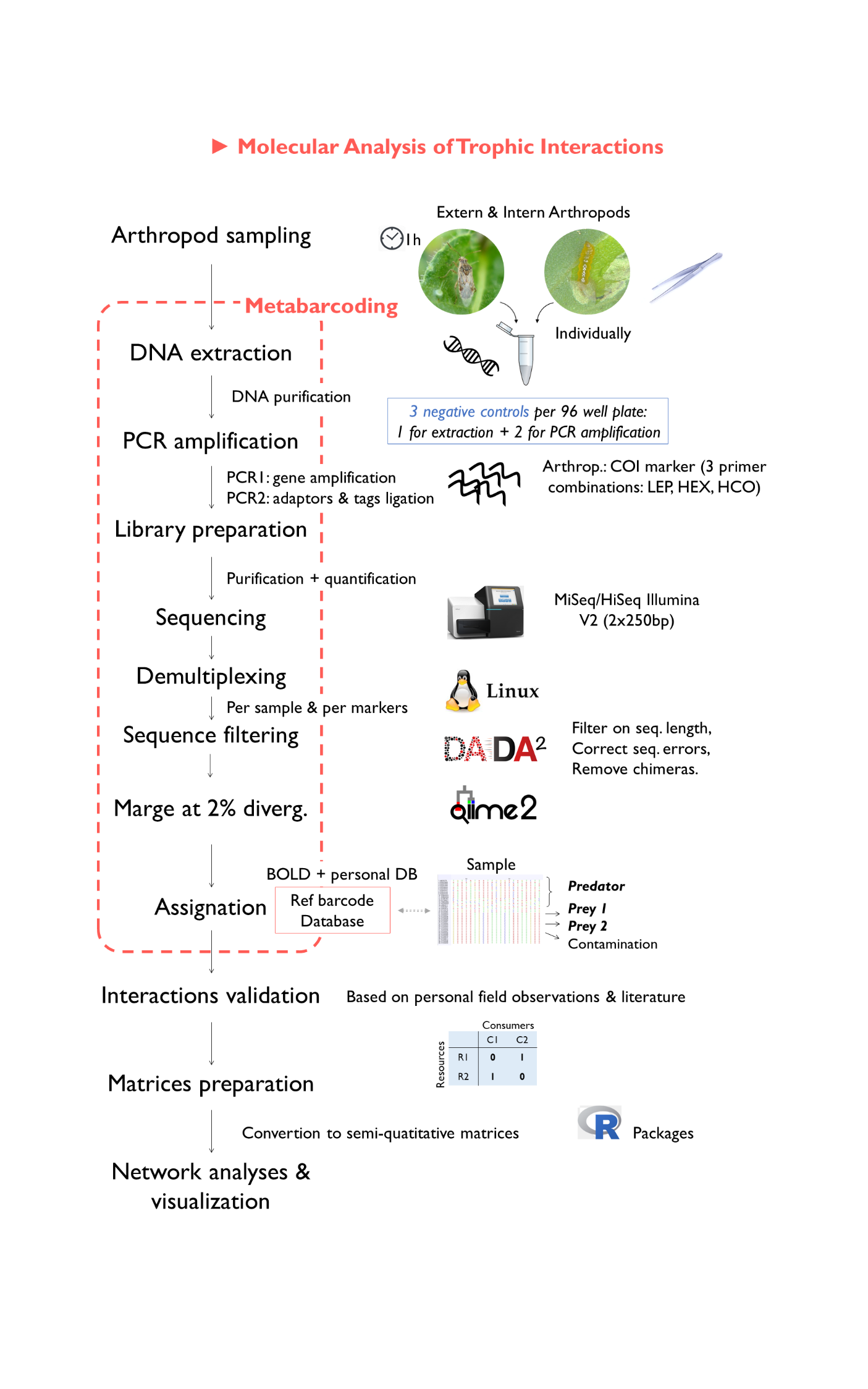

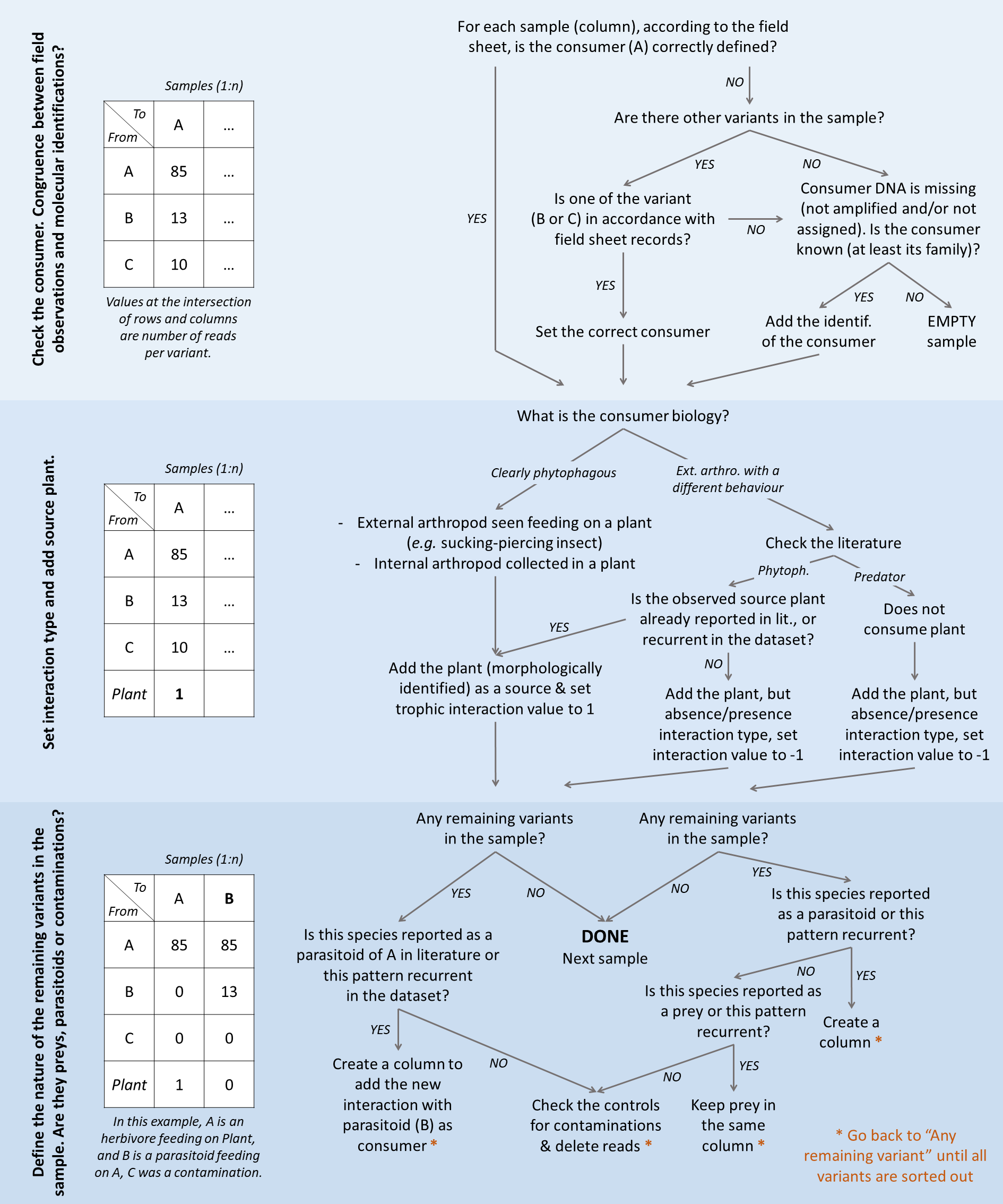

Figure S2: Flow diagram presenting the process followed for the transformation of raw data from sequencing into an adjacency matrix.

Figure S3: Venn diagram of taxonomic coverage obtained with the three COI markers used in this study, represented through a) arthropod family diversity and b) arthropod species diversity.

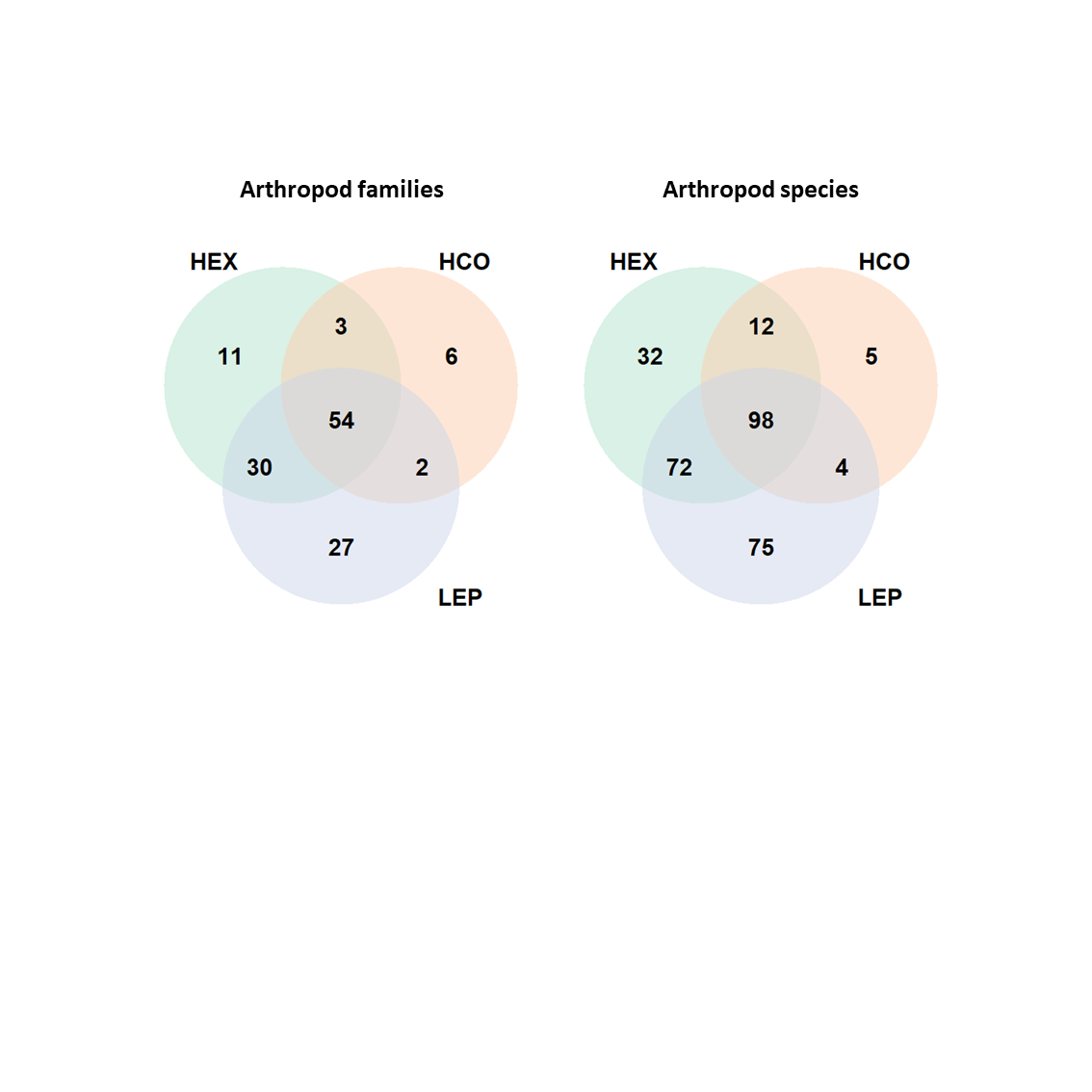

a)

b)

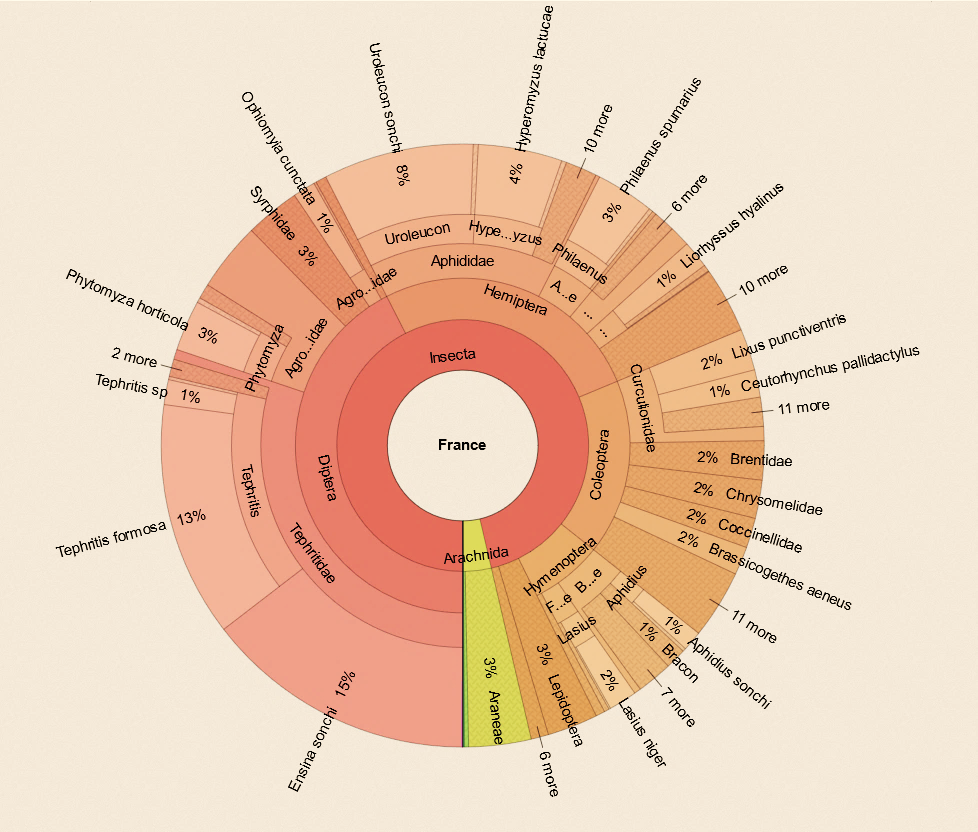

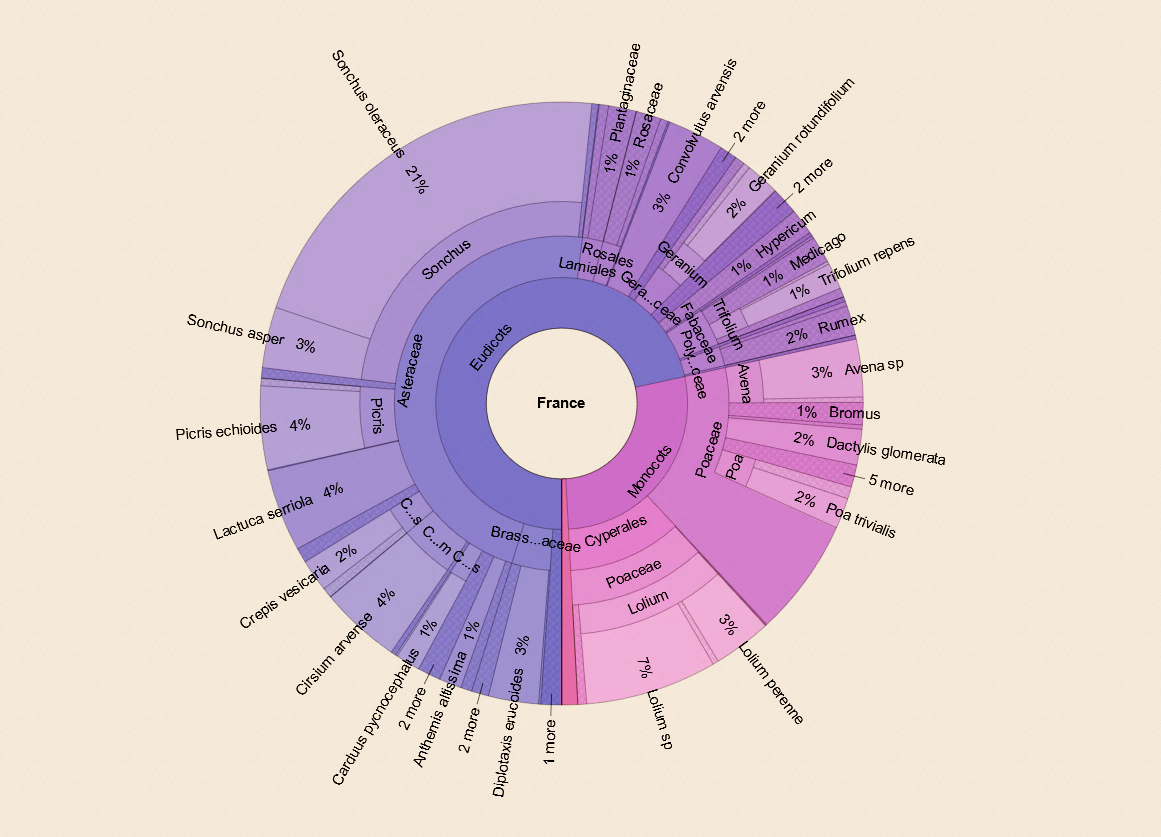

a)

b)

Figure S4: Taxonomic abundance of a) plants and b) arthropods in the dataset after data validation. This diagram allows visualizing the hierarchical organisation of taxonomic affiliation for each species, considering species abundance. E.g. Tephritis Formosa and Ensina sonchi (Tephritidae: Diptera) were two arthropod species very frequently sampled during this campaign (Spring 2018) representing 13% and 15% of samples collected, respectivelly.

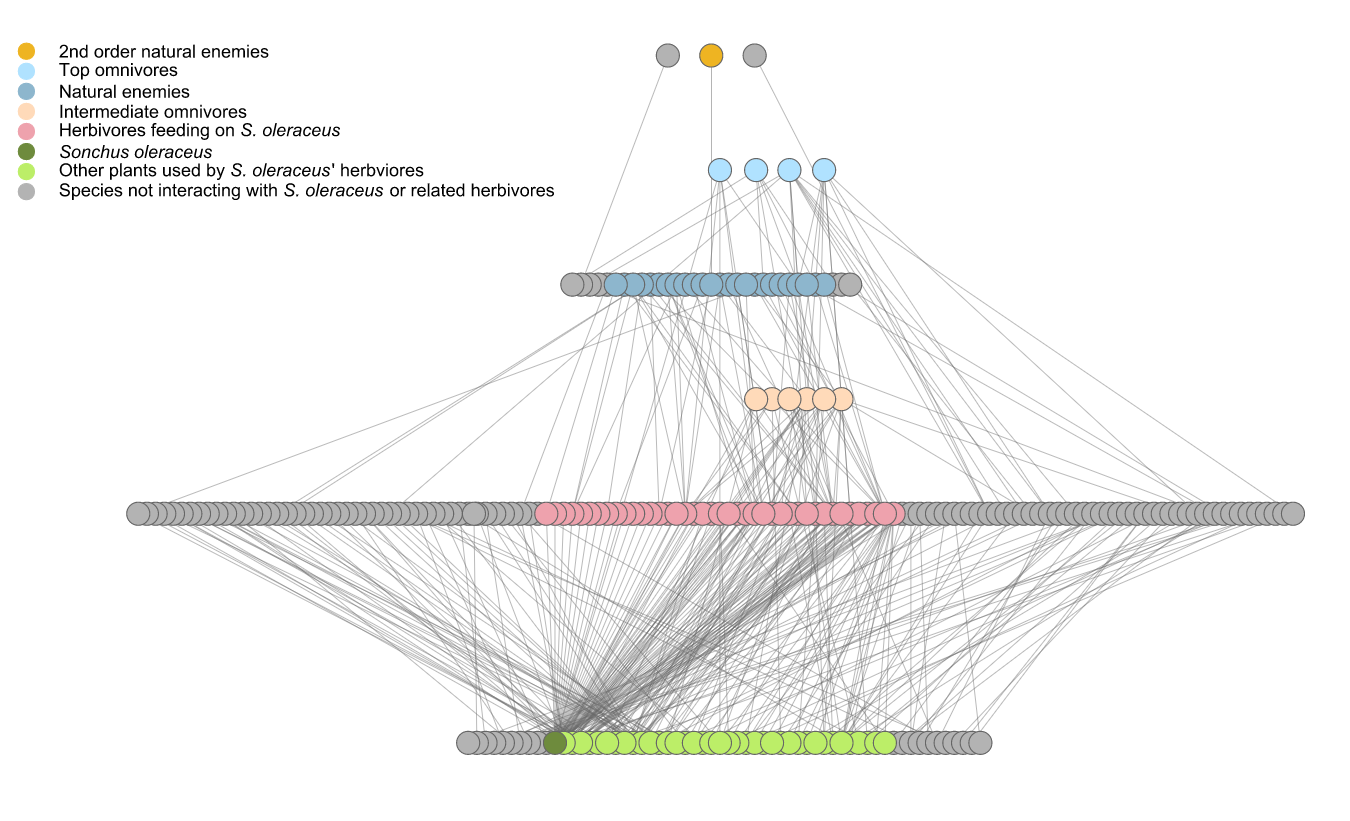

Figure S5 : Meta-network reconstructed through intense sampling in France during spring 2018. Collored nodes refer to subnetwork centred on S. oleraceus related interactions. The meta-network is composed of 241 nodes and 350 links, the subnetwork is composed of 116 nodes and 213 links.
